# Disentangling microbial associations from hidden environmental and technical factors via latent graphical models

**DOI:** 10.1101/2019.12.21.885889

**Authors:** Zachary D. Kurtz, Richard Bonneau, Christian L. Müller

## Abstract

Detecting community-wide statistical relationships from targeted amplicon-based and metagenomic profiling of microbes in their natural environment is an important step toward understanding the organization and function of these communities. We present a robust and computationally tractable latent graphical model inference scheme that allows simultaneous identification of parsimonious statistical relationships among microbial species and unobserved factors that influence the prevalence and variability of the abundance measurements. Our method comes with theoretical performance guarantees and is available within the SParse InversE Covariance estimation for Ecological ASsociation Inference (SPIEC-EASI) framework (‘SpiecEasi’ R-package). Using simulations, as well as a comprehensive collection of amplicon-based gut microbiome datasets, we illustrate the method’s ability to jointly identify compositional biases, latent factors that correlate with observed technical covariates, and robust statistical microbial associations that replicate across different gut microbial data sets.

---

High-throughput amplicon and metagenomic sequencing techniques are transforming our understanding of microbial ecosystems. Quantifying the relationships between microbes and their environments is at the core of microbial ecology, and sequencing has enabled quantitative surveys of microbial abundances in their natural habitats, providing microbial bio-geographical information at ecosystem scale [48, 23, 21, 13]. Targeted sequencing of marker genes, including bacterial 16S rRNA [45], eukaryotic 18S rRNA [47], and internal transcribed spacer (ITS) sequences [49] provides a proxy for microbial species abundances via read counts of operational taxonomic units (OTUs) [28] or amplicon sequence variants (ASVs) [6]. These data provide the means to measure not only microbial diversity and phylogeny in the environment but also statistical taxon-taxon and taxon-environment associations [17, 20, 30].

Statistical associations derived from sequencing data may not reflect direct ecological relationships, such as mutualism or competition. However, they can provide testable hypotheses about these interactions. One prominent example includes the discovery of a novel endosymbiotic relationship between the acoel flatworm *Symsagittifera sp*. and photosynthetic green microalgal, *Tetraselmis sp*., from co-occurrence patterns in the TARA ocean data [32]. However, technical limitations in experimental design and data collection procedures can confound the detection of reproducible and stable statistical relationships among the microbial constituents. Each step in an experimental workflow, including cell lysis, primer selection, PCR amplification, sample multiplexing, and sequencing platform can introduce biases [38]. Variability in library size [39] and batch effects [46] also influence the observed read counts. Moreover, the grouped taxonomic units are typically high-dimensional with respect to sample size, exhibit an excess amount of zeros, and carry only proportional (compositional) information [24]. Thus, dedicated methods are needed to estimate co-occurrence, correlation, and partial correlation for these data.

Biotic and abiotic factors have large effects on observed taxa compositions and co-occurrence patterns. For instance, environmental determinants, such as pH gradients, regulate soil microbial communities [31]. Host factors such as intra-family transmission [5], diet [14], antibiotic treatment [44], and disease states [11, 41] alter bacterial compositions in the mammalian gut. In light of the effects of experimental and biological factors on co-occurrences, it is challenging to disentangle direct microbial associations from those induced by unobserved latent covariates. If all relevant factors are known and measured, joint modeling of these covariates and microbial compositions can reduce the number of false positive taxon-taxon associations [3, 53]. However, for hypothesis-free population surveys, it is unlikely that all factors can be correctly specified *a priori*.

Here we introduce a parsimonious and computationally tractable procedure to decompose the matrix of statistical associations estimated from sequencing data into: (i) a sparse set of direct microbial associations as quantified by partial correlations among the microbial compositions, and (ii) a set of latent associations induced by hidden covariates (e.g. environmental and batch effects) and represented as “low-rank” components in the association matrix. This representation is known as latent variable graphical model in the statistical literature and can be estimated efficiently via convex optimization [10]. We show, in theory and practice, that the proposed model captures both compositional and unobserved latent effects and allows a parsimonious and consistent representation of estimated taxon-taxon associations. Using a compendium of gut microbial datasets, we demonstrate that several estimated latent factors strongly correlate with observed technical covariates. We detect a core set of statistical taxon-taxon associations across these datasets, which are amenable to follow-up experimental validation.

## Results

### A latent variable graphical model for microbial associations

Our modeling approach starts with a given microbiome data set *Z*, derived from targeted amplicon or metagenomic sequencing. The matrix 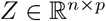 comprises *n* rows, representing the samples, and *p* columns, representing the measured taxa (e.g., OTUs, ASVs, or miTAG OTUs [35]). Each entry *Z_ij_* is a read count, absolute or relative abundance, presence/absence, or a transformed quantity representing the taxon. Our goal is to estimate pairwise associations between taxa via an appropriate measure of association. The collection of non-zero or statistically significant associations can then be summarized as a taxon-taxon association network.

A popular linear measure of association is covariance or, when data are standardized, Pearson correlation. In the microbiome context, covariance estimators are available both for compositional data [20, 8] and for absolute microbial abundance data [54]. To account for transitive correlations, estimators of the inverse covariance (or precision) matrix are also available [30, 16]. The inverse covariance matrix encodes the *conditional* dependency structure of the underlying variables and provides a parsimonious summary of (direct) taxon-taxon associations. In practice, the number of taxa often exceeds the number of samples in microbial datasets, thus requiring regularized estimators of the inverse covariance matrix. Given a *p* × *p*-dimensional empirical covariance (or correlation) matrix 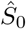, a common estimation framework is based on solving a convex optimization problem of the form:

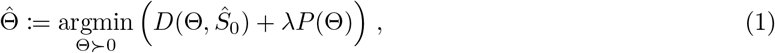

where 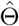 is the desired inverse covariance estimate, *D*(·, ·) a log-likelihood data fitting term, *P*(·) is a penalty function that enforces structural assumptions about the estimates, and λ > 0 is a scalar tuning parameter that balances both terms. The positive definiteness constraint Θ ≻ 0 ensures that 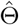 is invertible.

The log-likelihood function *D*(·, ·) is often taken from the exponential family of models (e.g., Gaussian, Poisson or binomial distributions) and reflects the assumptions on the data-generating process underlying *Z* [43, 52]. A popular choice is the log-determinant divergence 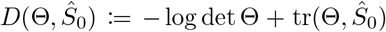 which encompasses several exponential family models [43, 27].

A common penalty function that allows efficient statistical estimation and parsimony is the choice *P*(·) = ║·║_1_, the entry-wise ℓ^1^ norm [40, 19, 27, 55]. The resulting estimates 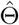 are generally sparse, and the non-zero entries in 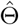 can be interpreted as direct statistical taxon-taxon associations.

Estimators of this type implicitly assume that all variability in the data can be explained by the covariation of all measured taxa, and with only a few conditional dependencies per taxa. However, as environmental covariates, latent functional mechanisms, and technical artifacts can influence taxa-taxa associations, sparse estimators, including SPIEC-EASI [30], cannot capture the effects that give rise to the corresponding inverse covariance patterns. This requires a more general notion of parsimony through the concept of latent variables. Here, the inverse covariance matrix is assumed to comprise two parts Θ ≔ *S* – *L* where the matrix *S* is sparse and represents direct associations between the taxa and the matrix *L* is dense but of “low rank”, i.e., it represents a superposition of outer products of few latent variable vectors [10]. Such a “sparse and low-rank” (slr) model is promoted via the following penalty function:

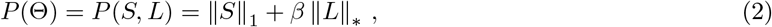

where the ℓ^1^ norm encourages sparsity in *S*, the nuclear norm ║·║_*_ provides a convex approximation to the rank, and the parameter *β* ≥ 0 regulates the importance of the latent variable components in *L*.

We illustrate the idea of the latent graphical model with a toy example, shown in Figure 1. Assume that the variability of *p* =16 taxa is characterized by a network of 10 dependencies. Additionally, the first five taxa rely on an (unknown) shared nutrient but do not interact with each other. In this setting, the population inverse covariance matrix of this system takes a form as shown in the left panel of Figure 1. If the shared nutrient influence is not taken into account, the dense 5 × 5 block of negative inverse covariance entries could be interpreted as tightly coupled direct association network among the five taxa. Moreover, given only relative abundance data, every element of the inverse covariance matrix is obfuscated by nonzero values due to the compositionality of the underlying data (as sketched in Figure 1, left panel). Sparse inverse covariance estimators can reduce, but not eliminate, the number of spurious edges in this network (as exemplified in Figure 1, bottom).

**Figure 1:**
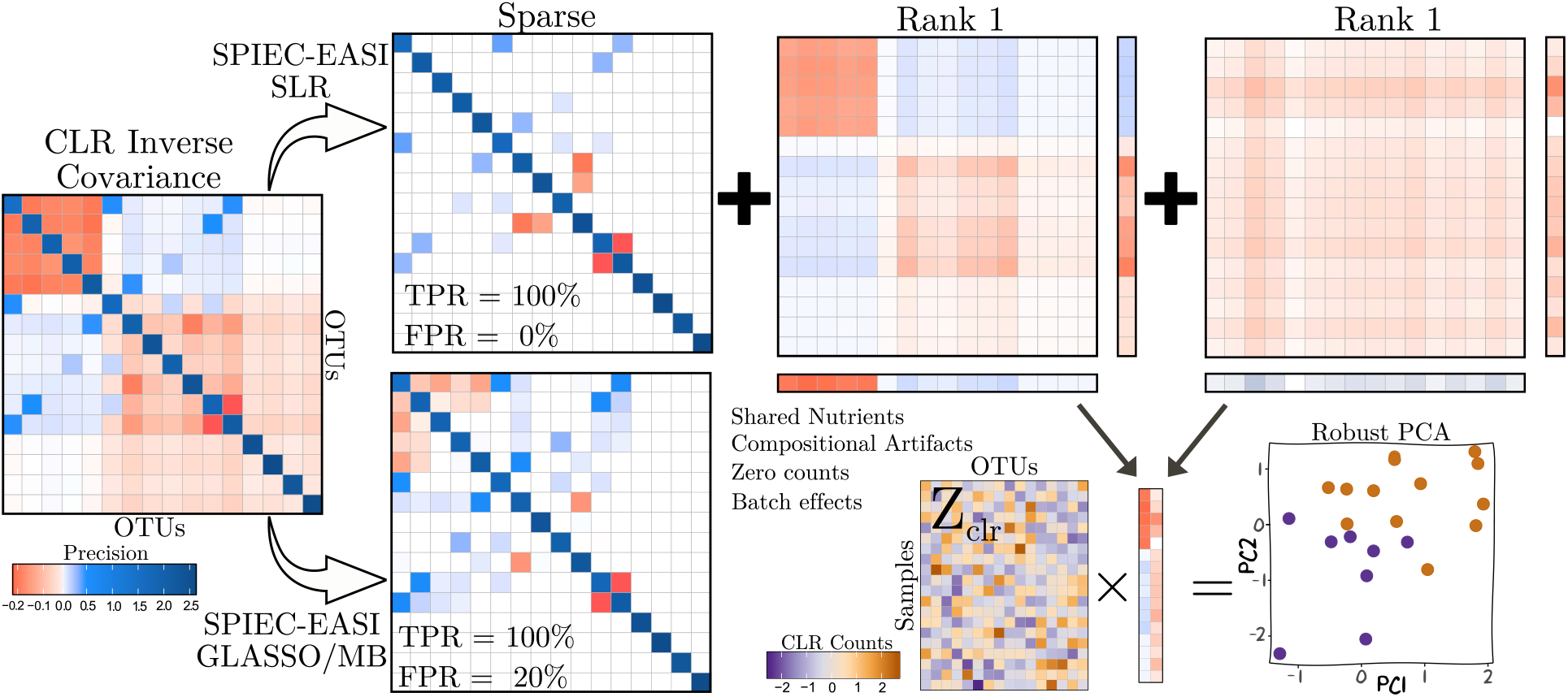
Illustration of the latent graphical modeling approach on a toy example. Left: Observed population inverse covariance matrix. Top: Latent graphical model decomposition (as implemented in the SPIEC-EASI slr framework). The model learns the correct set of direct associations (top left matrix) and two latent factors, corresponding to environmental (top middle) and compositional effects (top right). The latent factors (marginal vectors of the two matrices) can serve as robust principal components for lowdimensional representation of the sample space (bottom right). Bottom left: SPIEC-EASI with sparsity assumption (glasso or mb) misidentifies associations from latent effects as direct associations (red entries in top corner).

Partitioning covariations into those coming from direct associations and those coming from hidden factors can be particularly effective when the latent covariations are spread out over all entries of the inverse covariance matrix or when similar covariation patterns are concentrated over a large-enough subset of variables.

Our toy example presents both these scenarios. The upper panels in Figure 1 show how the observed inverse covariance matrix can be decomposed into three distinct parts: a sparse matrix capturing the direct associations, a dense block-structured matrix of rank one, capturing the shared nutrient factor, and a dense hub-like matrix of rank one, capturing compositional artifacts. This leads to a parsimonious description of the inverse covariance with the ten correct direct association entries, the *S* matrix, and two *p*-dimensional latent vectors (as shown by the marginal vectors in the upper right two panels), forming the *L* matrix. The collection of latent vectors also allows the construction of robust principal components (PCs) [7] which facilitate a low-dimensional description of the sample space (see bottom right panel of Figure 1). The scores of the robust PCs may correlate with measured meta-data such as, in the toy example, batches where the unknown co-varying nutrient is at different levels (scatter plot, bottom right of Figure 1).

Combining the log-determinant divergence with the penalization in Eq. (2) leads to an estimator that can provably identify the sparse and the low-rank part of Θ under certain model assumptions [10]. When latent effects only stem from total sum normalization, we provide novel theoretical recovery guarantees for compositional data in the Appendix. We solve the associated convex optimization problem in Eq. (1) using a variant of the Alternating Direction Methods of Multipliers (ADMM) algorithm proposed in [36].

### Computational inference framework

The latent graphical model is integrated into a comprehensive computational framework for association network inference from microbial count data, the R package `SpiecEasi` available at https://github.com/zdk123/SpiecEasi. SPIEC-EASI (SE) enables the use of a variety of graphical model inference schemes which all rely on specific combinations of data transformations, graphical models, optimization, and model selection.

Here we consider several instances of the framework. The latent, or sparse and low-rank (slr) graphical model, introduced here, is denoted by SE-slr and operates on centered log-ratio transformed count data [1]. The objective function in Eq. (1) with the penalty in Eq. (2) is optimized with ADMM [36] for a given pair of tuning parameters λ and *β*.

When the *β* = 0, i.e, no latent factors are assumed, the model reduces to the standard sparse graphical model. In SPIEC-EASI, two variants are available: the “graphical Lasso” [19] (SE-glasso) and its approximation via the Meinshausen-Bühlmann neighborhood selection scheme (SE-mb) [40]), as implemented in `huge` package [33].

For comparison, we also consider two novel SPIEC-EASI variants, SE-poisson and SE-ising. In SE-poisson, we first apply a common sum scaling [39], which divides each count vector by the smallest sample sum. This accounts for compositional effects by creating samples with identical count totals. In SE-ising, the count data were transformed into presence/absence data, i.e., binary data. Both variants then perform neighborhood selection using the generalized Lasso under a Poisson or Binomial link, respectively, as implemented in the `glmnet` R package [18]. We refer to the Methods section for further details.

In practical applications, the SE models require proper data-driven tuning of the regularization parameters. For the sparsity parameter λ, we use stability-based model selection (StARS) [34] to determine λ across the regularization path using the `pulsar` R package [42]. Briefly, we solve the corresponding graphical model over random subsamples, identify all edge sets, and compute an edge variability index along the entire regularization path. We select the λ parameter at which the edge variability of the graph is below the default threshold 0.05.

The latent graphical model (SE-slr) requires the additional tuning of the parameter *β*. Rather than varying the *β* parameter, we directly consider the rank *r* of the latent variable component *L*. For each *r* ∈ [2, *r*_max_] along the “rank”-path with *r*_max_ ≪ *p*, we solve the SE-slr model and use the StARS routine along the λ-path. Across all equi-stable sparse networks, we select *r* that minimizes the extended Bayesian Information Criterion (BIC) [33] of the full inverse covariance model.

### Disentangling compositional effects

We first investigated the ability of the latent graphical model (SE-slr) to identify direct associations that are obfuscated by purely compositional effects. We compared its performance to SE-glasso and the composition-adjusted thresholding (COAT) estimator [8] under two different scenarios: sparse networks generated (i) from synthetic absolute abundance data and (ii) measured quantitative microbiome profiling (QMP) data [51].

#### Performance on synthetic data

We generated absolute abundance covariance matrices Ω induced by sparse inverse covariances Ω^−1^ with three different graph topologies: band, cluster, and scale free (see Figure 2B). We refer to [30] and the Methods for details). Each method receives as input the covariance matrix 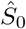 of the clr-transformed *relative* abundance data. The task is to recover the true network, i.e., the nonzero entries of Ω^−1^. We consider seven different network sizes *p* ∈ [16, …, 150] and report recovery results averaged across *R* = 10 replicates. For each scenario, we report each method’s empirical estimate that minimizes the relative Hamming distance (the sum of edge disagreements divided by number of edges in the true graph) over the entire solution set. Figure 2A summarizes the performance of the three methods (top row) as well as sparsity levels of the Ω, Ω^−1^, and the maximum degree *d*_max_ of the underlying network. We found that SE-slr achieved near-optimal recovery across all graph topologies, vastly outperforming SE-glasso and COAT. SE-glasso converges to SE-slr levels of performance at *p* = 150. The COAT estimator is outperformed in all scenarios, implying that there is no thresholded covariance that can recover the support of its sparse inverse.

**Figure 2:**
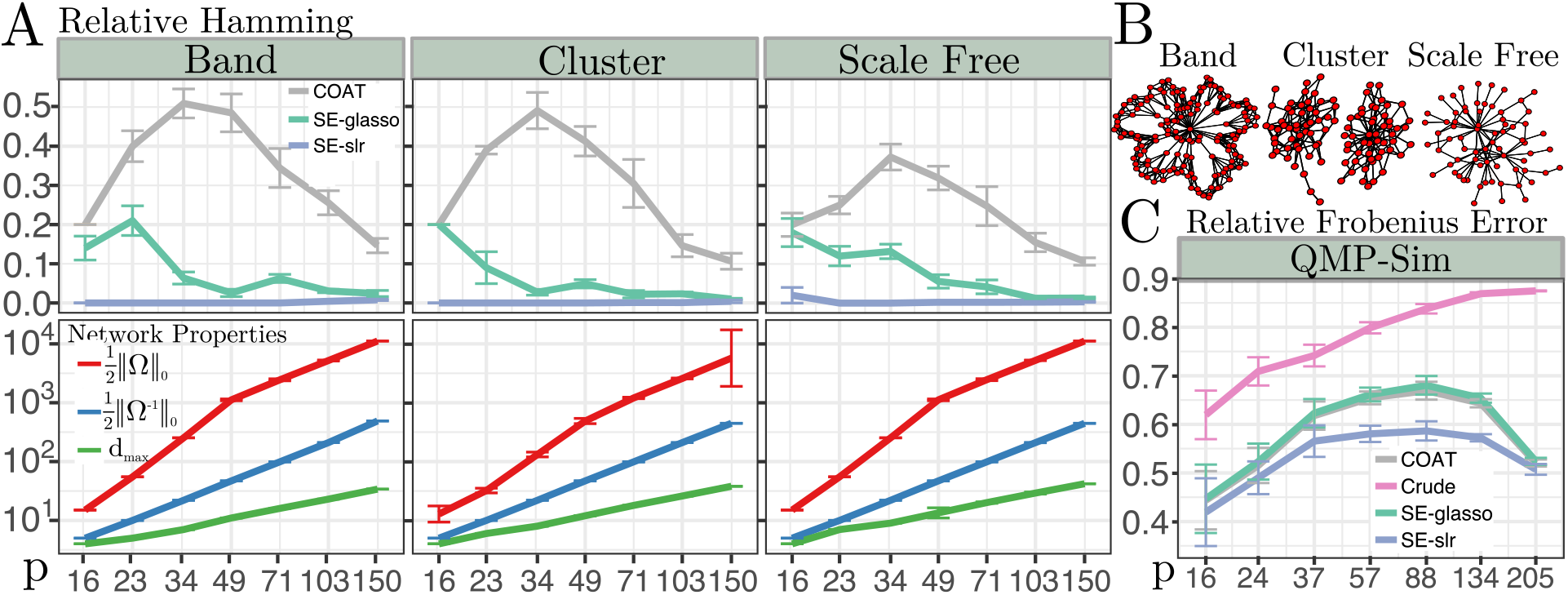
**A**: Asymptotic performance of COAT, SE-glasso, and SE-slr. Top: Hamming distance between the inferred and true graph. Bottom: Properties of the network and covariance matrices. **B**: Examples Band, Cluster and Scale-free graphs, used in simulations. **C**: Performance of simulated Quantitative microbiome profiling (QMP) simulations in relative Frobenius error norm of inverse covariance estimators, and the raw compositional covariance.

#### Covariance matrix recovery on QMP data

Recent experimental advances allow the joint quantification of total bacterial loads and relative abundances of OTUs in fecal samples [51]. These data provide an empirical basis to compare covariance estimates from compositional data sets to covariances of absolute abundance data. We used the profiling data from [51] to simulate QMP data (QMP-sim, see Methods), thus preserving the covariance structure from the absolute abundances. As the true inverse covariance matrix is not known, we evaluated each method’s performance by measuring relative Frobenius norm between the log-transformed QMP-sim covariance matrix and the sample covariance from untransformed compositions, the COAT estimator, and the *covariance matrices* 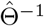, derived from SE-slr and SE-glasso, respectively. We again report the “oracle” performance of each estimator in Figure 2C. Covariance estimates from SE-slr outperformed all other estimates. SE-glasso and COAT’s performance was comparable and provided substantial improvements over the “crude” compositional sample covariance estimator. At *p* = 205, SE-slr, SE-glasso, and COAT achieved near-identical Frobenius norm.

### Consistent network estimation on American Gut Project data

We next investigated the ability of the latent graphical model to achieve consistent and reproducible microbial network inference in the finite sample setting using data from the American Gut Project (AGP) [37]. We first partitioned AGP into a large-sample reference set and sub-sampled OTU tables (see Methods). For each network inference method, we computed the “oracle” performance, averaged over ten replicate subsamples, in terms of relative Hamming distance between the inferred networks from subsamples and the respective large-sample reference network.

As shown in Figure 3A, the graphical models varied greatly in consistency even when thousands of samples were available. SE-slr showed dramatic consistency improvements compared to the other methods. Among previously published methods, SE-glasso showed the best consistency. SE-poisson’s relative Hamming distance exceeds 0.7 even when *n* = 3551 samples are used, making it particularly unsuited for reproducible microbial network inference.

**Figure 3:**
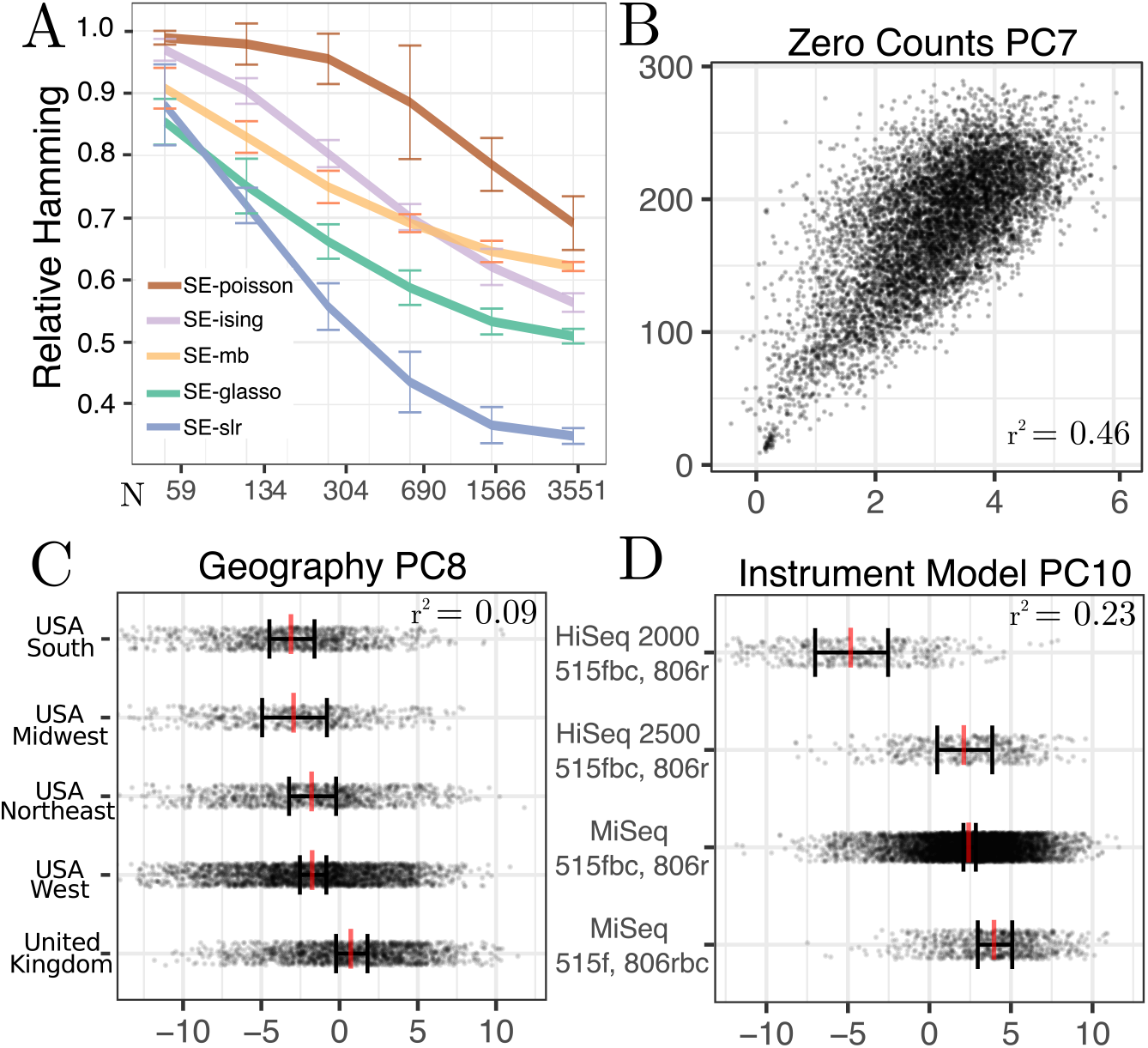
American Gut Project (AGP) microbiome **A:** We infer networks from subsampled test AGP rounds, compared to the network learned from all reference rounds. OTU tables are fixed at *p* = 306 and, AGP rounds are split into a test and reference sample subsets. Each method serves as its own reference, for which we compute the relative Hamming distance 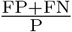, where FP is number of false positive edges, FN is number of false negative edges and P is the number of true edges in the reference network. **B, C, D**: We constructed the robust PCA from the low rank matrix of the SE-slr selected inverse covariance components, and found meta-data that correlated with **B:** PC7 (zero counts per sample, *r*^2^ = 0.46), **C:** PC8 (geography, *r*^2^ = .09) and PC10 (instrument model, *r*^2^ = 0.23).

To elucidate factors that contributed to improvements in SE-slr consistency, we correlated the robust PCs derived from SE-slr’s *L* matrix with covariate data available in the AGP (Figure 3B-D). We observed that the latent components PC7 and PC10 strongly correlated with sample features, including the number of zero counts in the sample (Figure 3B) and sequencing platform (Figure 3D). We also observed that PC8 correlated modestly with geographic origin of the subject (Figure 3C).

### Gut microbiome network meta-analysis

Removing the influence of latent factors from network inference also enabled us to investigating the consistency of statistical microbial associations *across* publicly available human gut microbiome studies. To conduct such a meta-analysis on a large scale, we searched the QIITA database (https://qiita.ucsd.edu, [25]) and compiled a comprehensive collection of human gut microbiome 16S rRNA sequencing data sets, each consisting of at least 100 samples. The final collection of 26 distinct datasets (see Methods and Appendix for an overview and citations) comprised 3032 OTUs across 26301 samples. On each data set, we inferred microbial association networks using our five SPIEC-EASI variants and performed a meta-analysis of the derived networks and latent components.

#### Network consistency across datasets and methods

To assess the consistency of microbial associations across the 26 datasets we first identified, for each pair of datasets, the set of common OTUs between them. This resulted in 215 pairs of OTU sets of sufficient size (*p* > 40). For each inference method, we then compared the similarity between the induced sub-networks by measuring the fractions of sign-consistent edges between corresponding OTUs and transforming these fractions into normalized “edge distances” between networks, see Methods Eq. (11). We found (see Figure 4A) that SE-slr networks have the lowest median edge distance between datasets (SE-slr: 0.565, SE-gl: 0.611, SE-ising: 0.716, SE-poisson: 0.769). Among the 215 cross-dataset network comparisons, the SE-slr method achieved the smallest edge distances for 121 pairs (> 55%), followed by SE-glasso which is minimal for 66 pairs (≈ 30%). All other methods produced ≤ 11 minimum edge distance network pairs.

**Figure 4:**
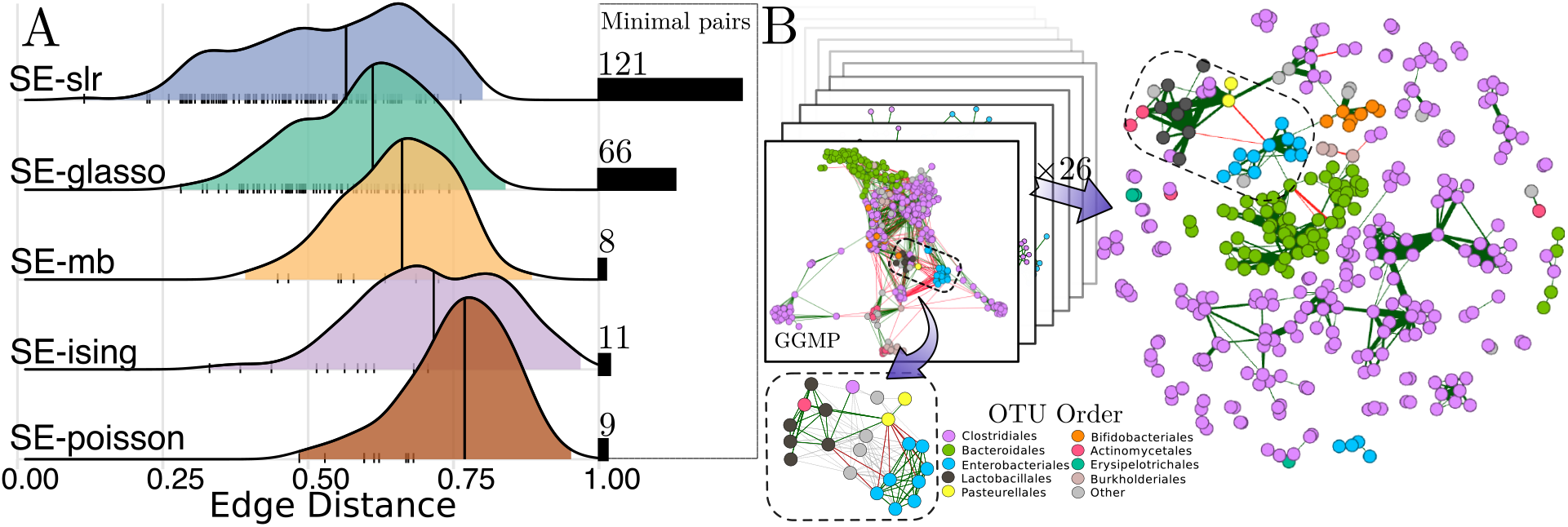
**A:** Pairwise comparisons of the networks inferred among 26 human gut microbiome data sets obtained from the QIITA database. Edge distance is derived from the proportion of shared edges in subgraphs from the set of common OTUs. Distribution of edge distances for each method, with tick marks indicating the minimum observed distance for a particular pair of datasets. Horizontal bars show the number of pairs the corresponding method achieved the minimal edge distance. **B:** Top right: The SE-slr consensus network across 26 human gut microbiome datasets (schematic on left). Node color indicates OTU taxonomy at the order level, edge color indicates consensus positive (green) and negative (red) association. One particular module in the consensus network is highlighted (dashed box), together with that module in the Guandong Microbiome Project(GGMP) network (bottom left, see text for details).

#### Robust PCA of latent components

We next analyzed the properties of the collection of latent components inferred with the SE-slr model across all 26 datasets. For each dataset, we first constructed the robust principal components by orthogonalizing the latent factors *L*. In the absence of known latent factors, we correlated the resulting principle components with experimental meta-data associated with each dataset. We classified the available sample meta-data into four broad categories: I) subject-specific features, including sex and age, II) technical features, such as primer plates and run prefix, III) sequencing characteristics, including sequencing depth and number of taxa with zero counts, and IV) host life style, including diet and antibiotics use. We found that the number of latent components detected in the SE-slr models across all 26 datasets was correlated with the proportion of statistically significant meta-data variables (Spearman *ρ* = 0.48, Figure 5A), using correlation tests (for continuous variables) or ANOVA (for discrete variables), based on Bonferroni-corrected p-values at *α* =10 × –14. Further, we found that several covariates were consistently associated with latent effects with a large average effect sizes across multiple datasets (see Table 2). These corresponded primarily to batch effects and technical artifacts (e.g. number of OTUs with zero count, collection/run date and primer prep plate, Figure 5B/C, category III) as well as subject-level, biological and behavioral effects (subject id, age, sex, diet and antibiotic exposure, Figure 5B/C, category IV). Overall, the distribution of significant covariates ranged over all categories and possible effect sizes (Figure 5B), with `host_subject_id` being a particularly strong “biological” effect (Figure 5C, category I/red) and number of taxa with zero counts `zero_tax` a strong “sample sequencing” feature (Figure 5C, category III/green).

**Figure 5:**
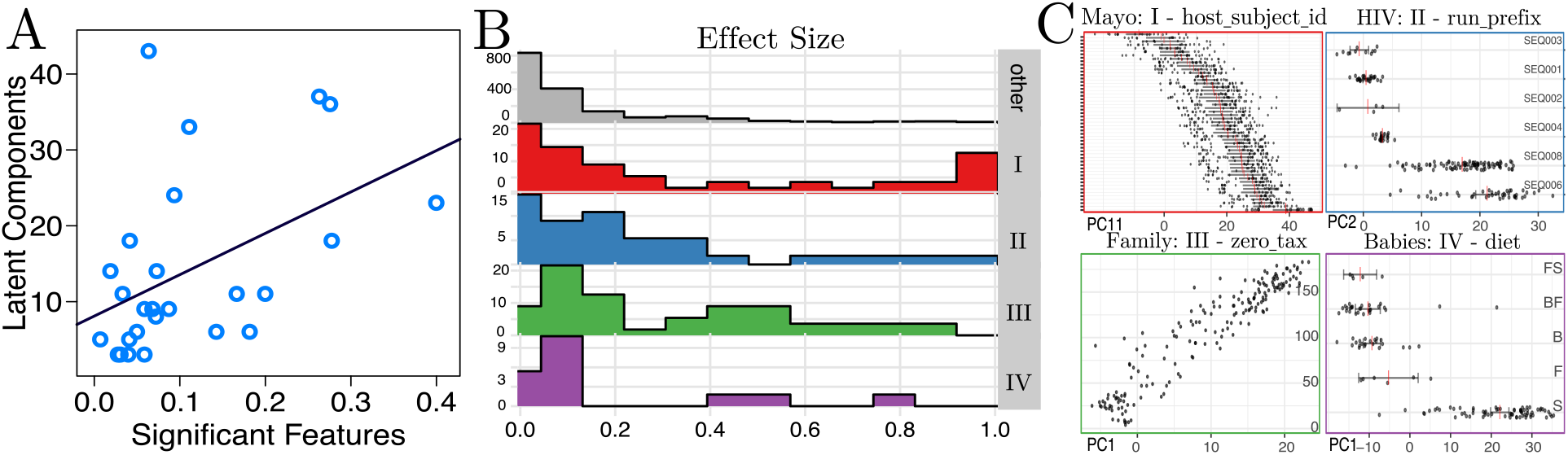
**A:** The number of latent components across 26 human gut SE-slr networks is correlated with the proportion of meta-data features that are significantly associated with scores of a robust PCA, based on Bonferroni-adjusted p-values at *α* = 10^−14^. **B:** The distribution of ANOVA or correlation test effect sizes in meta-data feature - robust PCA scores models, across all datasets. Meta-data feature categories are divided into I: `host_subject_id`, age, and `sex`, II: `run_prefix collection_timestamp`, `primer_plate, run_date`, and `extraction_robot`, III: `zero_tax` and depth, IV: `diet` and `antibiotics`. **C:** Example principal component scores vs a correlated meta-data feature for each of the above categories.

#### Consensus Gut Microbiome Network

Our network inference framework also allows for the construction of gut microbiome consensus networks, including edges across all datasets. Because of the improved consistency of the SE-slr model, we focused here on the analysis of the SE-slr consensus gut microbiome association network. From the 26 individual SE-slr networks, we identified and combined edges that were present in at least 20% of the datasets (see Methods). This resulted in a consensus network with 347 OTUs and 560 edges, shown in Figure 4B. This SE-slr consensus network analysis resulted in 44 network modules. While most modules consisted of few OTUs and/or were dominated by a particular taxonomic order, we identified one module with diverse lineages (highlighted in dashed box in Figure 4B), including members of Enterobacteriales, Pasteurellales, Clostridiales, Lactobacillales, and Actinomycetales. We found two negative edges in this network module that connected two subcomponents: one between OTU 4448331 (*Enterobacteriaceae sp*.) and OTU 4477696 (*Haemophilus parainfluenzae*), and one between OTU 4448331 and OTU 4425214 (*Streptococcus sp*.). These associations were particularly interesting because they were found in an independent experimental study [22] and were exclusively present in the SE-slr consensus network.

To highlight this unique ability of the SE-slr model, we extracted the SE-slr network module for one particular dataset, the Guandong Microbiome project (GGMP), and jointly visualized the direct associations in the GGMP network (colored nodes nodes and edges in Figure 4B outset) and the four latent components and OTU correlates (gray nodes and edges in Figure 4B outset). We observed strong positive correlations between the OTUs and the inferred latent variables. Thus, the negative direct associations could be detected only in the presence of the low-rank components. All other SE methods were unable to identify these negative associations.

## Discussion

Microbial association network inference has become a standard exploratory data analysis tool that enables the formulation of data-driven parsimonious hypotheses about the structure of microbial communities [2, 50, 44, 15]. However, the detection of microbial associations from high-throughput sequencing data is often obfuscated by experimental artifacts and noisy data generation processes. Due to both the difficulties discussed and the lack of gold standards in form of curated or experimentally-confirmed microbe-microbe interactions, the usefulness, validity, and applicability of network inference tools continues to be actively debated and researched [12, 9].

Here we present a computational tool that can handle a class of artifacts previously unaccounted for in microbial network inference procedures. While existing methods allow for measured covariates to be included as network nodes [3, 53, 4] or account for data compositionality [30, 8, 15], the latent graphical model, as implemented in the sparse and low rank SPIEC-EASI model (SE-slr), can incorporate arbitrary latent effects in microbial association network learning. This disentangles inference of a sparse set of direct associations from large low-rank sample-wide variations, induced by compositional, environmental, genomic-technology, and batch effects. SE-slr uses joint stability-based and BIC model selection to select a low-rank and a sparse inverse covariance matrix both of which offer valuable data summary capabilities. The low-rank matrix allows a robust low-dimensional description of the data via its principal components [7]. The sparse matrix can be interpreted as network of microbe-microbe associations which provide testable hypotheses about true microbial interactions in the ecosystem.

We have benchmarked SE-slr alongside other methods, including novel SPIEC-EASI variants and the COAT estimator [8], using synthetic data, quantitative microbiome data [51], and a curated collection of 26 gut microbiome data set, derived from the QIITA database (https://qiita.ucsd.edu, [25]).

The synthetic data simulations, both in the asymptotic regime and with finite QMP data, have been used to validate the theoretical guarantees we have derived for the SE-slr estimator on compositional data (see Appendix for mathematical details). We have shown that, given a sufficiently sparse population inverse covariance matrix, SE-slr can outperform the COAT estimator which also comes with statistical guarantees in the compositional setting [8] (see Figure 2).

Our analysis on the largest of the 26 gut microbiome datasets, the American Gut Project (AGP) data [37], has been instrumental in showing the superior consistency of SE-slr compared to other graphical models. By constructing a large sample reference network for each method, we can show that SE-slr converges considerably faster to its reference network than all other graphical models (Figure 3A). A comparison of the jointly derived robust principal components to available covariates in AGP identified several batch effects that likely hinder other graphical models, especially with regard to consistency. Important features include the level of excess zeros in the samples as well as the sequencing platform used (Figure 3B-D). The strong influence of the number of zeros in the samples may be alleviated by improved scale estimations prior to data transformations [29] or zero-aware rank-based transformations [54]. However, we also recognized that many inferred latent components for the AGP data did not correlate with any of the measured covariates. This implies that, while some of the observed network inconsistencies can indeed be apportioned to measured covariates in the AGP, major drivers of network variability are likely due to unmeasured latent factors.

Our meta-analysis of the complete set of 26 gut microbiome datasets revealed that the SE-slr model also allows more consistent network estimation *across* independently inferred networks (see Figure 4A). This allowed the construction of a gut microbiome consensus network whose associations are conserved *across* different populations and experimental setups. While most associations in the consensus network are positive and link taxonomically related OTUs, we identified one heterogeneous network module that primarily represents residents of the upper digestive tract [26]. There, we identified negative associations between *Enterobacteriaceae sp*. and *Haemophilus parainfluenzae* and *Streptococcus sp*. that were conserved across at least five SE-slr networks but completely absent in consensus networks derived from the other methods. Interestingly, these same facultative anaerobes were described as conditionally rare taxa in gut microbiome time series; undetected at most time-points and with shared autoregressive structure driven by host niche and diet [22]. Although we do not explicitly model time dependency in our graphical model, we hypothesis that learning the latent components is sufficient to detect negative associations that are otherwise obfuscated by host effects. This analysis exemplifies that the large set of conserved microbial associations derived here may be indeed a good candidate list for true gut microbial interactions.

Going forward, we envision that the latent graphical model, as implemented in SE-slr, will become instrumental in reproducible microbial association detection from large-scale sequencing data, and ultimately, help unravel the complex interactions in microbial ecosystems in their natural habitats.

## Methods

### Data processing and OTU count transformations

We are given OTU count data in matrix 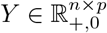, where 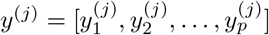 denotes a *p*-dimensional row vector from the *j^th^* sample, with total sample counts 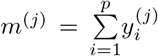. We assume that these counts are drawn from unobserved basis count data 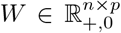 with 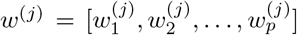 and with total community size 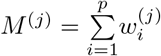.

We normalize the raw count data *y*^(*j*)^ with respect to the total count *m*^(*j*)^ of the sample. This gives us a vector of relative abundances - or taxa proportions/compositions - 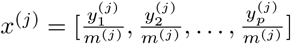.

#### Log-ratio transformations

Rather than considering raw compositions, log-ratios of the form 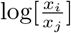 provide a rigorous basis for analyzing compositional data. The simple equivalence

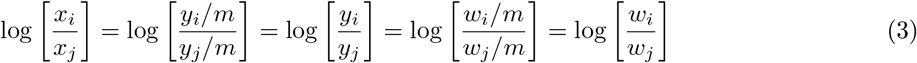

implies that statistical inferences drawn from the analysis of log-ratios of compositions are equivalent to those drawn from the log-ratios of the unobserved, absolute count basis. Several different log-ratio transformations have been proposed for compositional data [1, 69] and for microbiome data [81]. Here, we consider the centered log-ratio (clr) transform

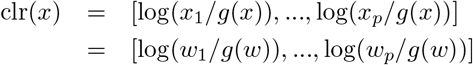

where 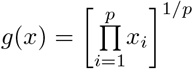 is the geometric mean. We collect transformed taxa compositions into a data matrix 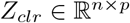.

#### Scaled Counts

An alternative to log-ratios is to use the count matrix *Y* directly. For network inference, this has been proposed in [3]. To control for differences in sequencing counts on standard normalization is the common-scale normalization [39]. Here, a count vector from sample *j* is given by 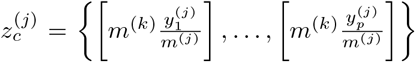, where *m*^(*k*)^ is the total count size for sample *k*, the sample with the lowest depth, and [·] rounds the proportion to the nearest integer. The sample space of *Z_c_* remains 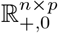.

#### Simple Presence/Absence transformations

The most radical approach to normalization is to remove count information entirely, and consider only the observed occurrences of an OTU in a sample. This leads to binary data via 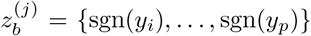, where sgn is the standard signum function, 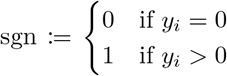, which covers all cases for non-negative count data. Binary data is stored in the matrix *Z_b_* ∈ [0,1]^*n*×*p*^. This data can then be used to detect co-occurrence patterns.

### Undirected graphical models from exponential families

Graphical models, or Markov random fields, describe conditional independence between nodes in a graph and can thus account for transitive correlations between nodes that are not directly connected. Formally, a graphical model is a family of probability distributions that factorizes according to the structure of a undirected graph 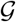. The graph 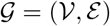 comprises the sets of nodes 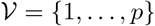 and (undirected) edges 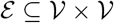 where 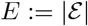 denotes the number of edges in the graph.

We denote by *e_ij_* (or *e_ji_*) the edge between node pair (*i*, *j*). In a graphical model, each vertex 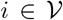 is associated with a random variable *z_i_* taking values in a space 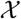, and the *p*-dimensional vector *z* = (*z*_1_, *z*_2_, …, *z_p_*)^⊤^ is Markov with respect to 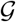. This means that random variables *z_i_* and *z_j_* are conditionally independent given *z_V_*, i.e., 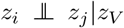, with *V* ∈ {1, …, *p*} \ {*i*, *j*} if and only if 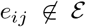. For nodes *V* ∈ {1, …, *p*} \ *i*, node *j* is in *N*(*i*), the neighborhood of node *i*, if and only if 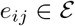.

Given *n* independent realizations of *z*, a key challenge in high-dimensional statistics is to derive and analyze consistent estimators of 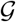 that are efficiently computable and are applicable in the *p* > *n* regime.

Conditional relationships between vertex pairs are typically characterized by a member *P* from the exponential family of probability distributions. That is, 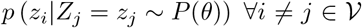, where canonical parameters *θ* = {(*θ_i_*, *θ_ij_*)}∀_*i*_ ≠ *j* are the intercept and conditional weights, respectively. We now briefly introduce the exponential graphical models. For extensive treatment, see references [85, 52].

The sufficient statistic for all distributions of interest here are linear functions of the data, so then we have the node-wise conditional distributions over *z*:

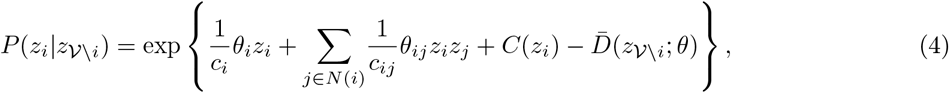

with positive constants *c_i_*, *c_ij_*, base measure *C*(*z_r_*) and log-normalization constant 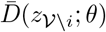. The joint distribution has the form

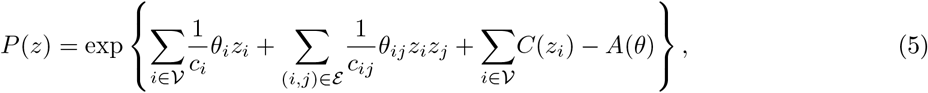

where *A*(*θ*) is the log-partition function that restricts *θ* to valid domains. Given continuous *Z_clr_*, count data *Z_c_*, or binary data *Z_b_*, introduced above, we can learn graphical models characterized by Gaussian, Poisson, and Ising models, respectively.

In the Gaussian case, we have that 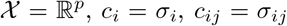 and 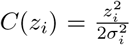. The values 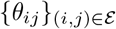 corresponds to the precision matrix, which must be positive definite.

For Poisson models, 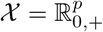, with *c_i_*, *c_ij_* = 1 and *C*(*z_i_*) = – log(*z_i_*!). For Ising models, 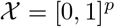 with *c_i_*, *c_ij_* = 1 and *C*(*z_i_*) = 0.

The goal of graphical model inference is to recover the graph structure, given *n* samples of a random vector, *Z* = [*z*^(1)^, …, *z*^(*n*)^] from a pairwise Markov random field, and specified by a sparsity-constrained conditional model. For the Poisson and Ising models, restriction of the domain makes learning a fully joint model difficult in practice, and then only for certain graph topologies [74, 85]. However, it has been shown that an estimate of the edge set can be constructed given 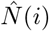 for each node: 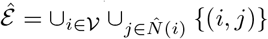.

Thus, we arrive at a general objective function for a regularized conditional log-likelihood to learn the conditional weight parameters for node *i*:

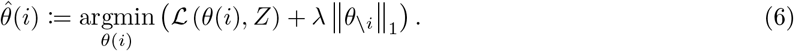

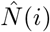 can be obtained from the non-zero pattern of 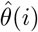. 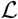 is the negative log-likelihood function associated with conditional distribution 4 and ║·║ is the vector ℓ^1^-norm, the sum of absolute values, parameterized by a positive λ. In the Gaussian case, the objective 6 corresponds to the neighborhood selection approach from Meinshausen and Bühlmann (2006) [40]. Efficient algorithms and theoretical recovery guarantees exist for Gaussian, Poisson, and Ising models [59, 58, 85]. The Poisson and the Ising model are now implemented in SPIEC-EASI (SE) and referred to as SE-poisson and SE-ising, respectively.

To learn a full, joint model for the Gaussian case, we have the objective

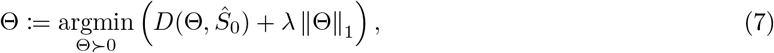

where 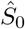 is the empirical sample covariance matrix, Θ is the positive definite precision matrix we wish to learn and ║·║_1_ is the sparsity-promoting matrix ℓ^1^ norm. *D* is the negative log-likelihood function of the multivariate Gaussian, 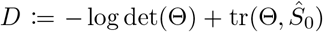. Objective (7) can be solved efficiently, for example, by the graphical LASSO (glasso) using a block-coordinate descent algorithm [19, 84].

### Latent variable graphical models via a sparse plus low rank decomposition

An important extension of Gaussian graphical model learning was proposed by Chandrasekaran et al [10] where, in addition to the *p* measurements (denoted by the set *O*), there are an additional *h* unobserved variables (set *H*). We have a Gaussian random vector 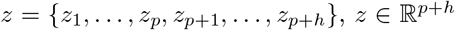. We have only access to the covariances of the observed variables, stored in the *p* × *p* matrix 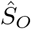. It is clear that the marginal inverse covariance matrix is not independent of the unobserved/latent variables, even in the sample limit, as

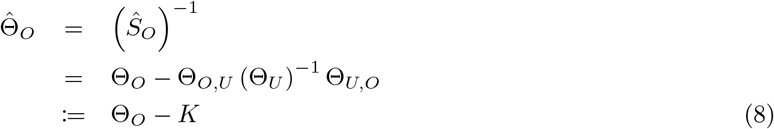

The second equality in 8 is given by the Schur complement of the partitioned precision matrix 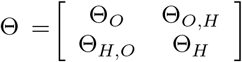. For real datasets, including microbial abundance data, there may be many latent effects. These cannot be easily conditioned out or eliminated by thresholding, unless all of the induced correlations are small. However, if *h* ≪ *p*, then *K* can be approximated by a matrix that has rank of at most *h*.

#### Latent compositional effects

For compositional data, in particular the clr-transformed compositions *Z_clr_*, a special instance of latent graphical model can be used to infer purely latent compositional effects when no unobserved dependencies or covariates are present. We denote the empirical clr-covariance matrix as 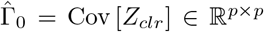 and log-basis covariance matrix as 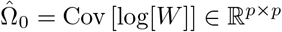. We define an inverse of the clr-covariance matrix

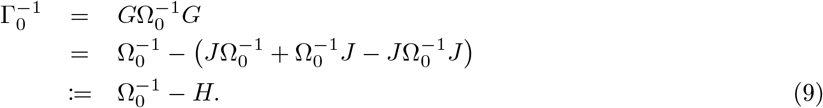

The matrix 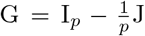 is the centering matrix, with *I* is the *p* × *p* identity matrix, 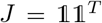 and 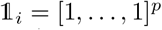 the *p*-dimensional all-ones vector. *H* is a symmetric, indefinite, rank-2 matrix, except when Ω^−1^ = *υI*, where rank(*H*) = 1.

Thus, the problem of identifying the basis inverse covariance matrix from compositional data is an special case of problem (8), to recover the components in a **sparse plus low rank** (slr) matrix decomposition [110]. Such problems are common in machine learning for matrix completion and robust PCA [61]. Because there are arbitrarily many decompositions of a matrix into two summands, unique identification of the model model requires the underlying generative model to follow certain conditions on the sparsity pattern and the low rank components.

Treating compositional *covariances* as a sparse plus low rank decompositions was first addressed in the composition-adjusted thresholding (COAT) estimator [64]. The goal of the COAT estimator is to identify the covariance matrix Ω rather than its inverse. COAT uses a one-step thresholding operation on 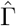. When *p* is large, this recovers Ω as long as 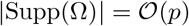 with non-zero entries bounded away from 0.

Here, we propose to explicitly model the rank-2 component *H*. We thus gain *exact* identifiability conditions from the rank-sparsity incoherence properties given by [63, 110, 10, 112] (see Appendix). Our general objective is to learn Θ as a proxy for Γ^−1^ as decomposition into a sparse matrix *S* and low rank components *L*

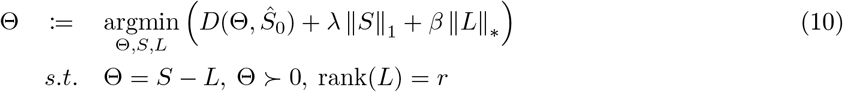

Objective (10) is similar to the sparse inverse covariance problem in objective (7), with additional equality constraints Θ = *S* – *L* and sparsity penalty on *S*. The nuclear matrix norm, 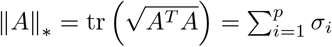, is a convex relaxation of the rank, and its minimization thus promotes low-rank properties of *L*. In the purely compositional setting, the parameter *β* can be tuned to achieve *r* = 2.

Objective (10) is nearly identical to the problems posed in [10, 36], except that we do not require the matrix *L* to be positive [semi]definite, thus allowing for negative eigenvalue in *H*. We use an alternating direction methods of multipliers (ADMM) algorithm to solve problem (10). Our R/C++ implementation is derived from the Matlab code to solve the primal problem, proposed in [36]. For fixed *β*, we use the Stability Approach for Regularization Selection (StARS) [34] to select an optimal λ along a user-provided path, as implemented in the pulsar package [42].

#### General latent effects

In the general case where both compositional and non-compositional latent covariates are present, we learn the rank of the matrix *L* by selecting the tuning parameter *β*, or equivalently, the rank *r*, via the Bayesian Information Criterion (BIC). This scheme is implemented and referred to as SE-slr.

### Constructing a robust PCA

The singular value decomposition of a symmetric *p* × *p* matrix *C* of rank *r* is given by 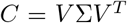, where *r* ≪ *p* and *V* is a *p* × *r* matrix. The principal component analysis (PCA) can be formed from the loading vectors 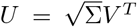. If we have a *n* × *p* data matrix *Z*, e.g. clr-transformed OTU compositions after mean-centering the columns, we form the “scores” of the PCA from the projection 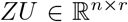, which are visualized via a scatter plot of a few columns of *ZU* [57].

For standard compositional data analysis, i.e., when *n* > *p*, we use 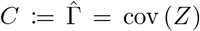, the empirical covariance matrix of *Z*, in which case *r* = *p* – 1. However, in real datasets (when *p* > *n*), the empirical covariance matrix will be rank deficient, and the SVD will not accurately represent the true components of the data variances.

We thus propose to set *C* ≔ *L*^−1^, the low-rank component of the inverse covariance matrix, inferred by SE-slr, yielding a robust PCA of rank 2 ≤ *r* ≪ *p* – 1 [7]. The SVD under consideration is *L* = *V*Σ^−1^*V^T^* with the corresponding loadings 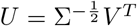. The *n* × *r* score matrix *ZU* thus corresponds to the projection of the data onto the principal components from the (inferred) marginal contribution of latent variables to the variance.

### Simulated compositional covariances

For our synthetic compositional covariance estimation experiments, we first created ten random replicates for cluster, band, and scale-free-type graphs each at node sizes of *p* ∈ [16,150]. For each graph, we set the number of edges to 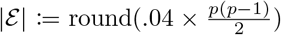 and the maximum degree to *d*_max_ ≔ max(2, ⌊*q* · *p*/4⌋), with *q* ∈ {0.9,1.0,1.1}, for the band, cluster, and scale-free graphs, respectively.

We randomly reassigned edges until the specified 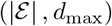 target is achieved, but protect edges whose removal would otherwise alter the number of connected components in the graph (bridge edges) [68].

From each graph, we generated a *p* × *p* basis precision matrix 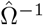 by assigning positive and negative weights to edges, drawn uniformly in [2, 3], and added a constant value to the diagonal to achieve a target matrix condition of 20. We converted this to the input CLR-covariance, 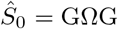, via a matrix inversion and the double centering transformation. This procedure coincides with the asymptotic performance for each estimator when the generating basis is a multivariate Gaussian.

### Quantitative microbiome profiling simulations

#### Data processing

We obtained 16S rRNA sequence data for 106 fecal samples from healthy subjects from the QMP, ERA study PRJEB21504, and bacterial load data from the supplement files of [51]. Using QIIME 1.9 [65], we clustered sequences to OTUs and assigned taxonomy via the sortmerna method under a 97% cluster similarity, and the closed-reference greengenes database. We merged common OTUs to the genus level using the tax_glom function in phyloseq R package [77]. This resulted in counts for 205 genera.

#### Model

Raw non-negative counts for each sample *j* ∈ {1, …, 106} were collected in a data vector 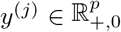. For each sample, we also obtained an estimate of the total community size, *M*^(*j*)^, given by the average of three flow cytometry-based measurements of the total bacterial loads from frozen fecal samples.

Our goal was to jointly estimate the OTUs proportions 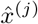, which has the standard unit-sum constraint 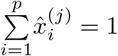, as well as the absolute abundance vector 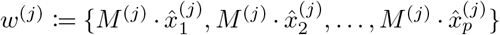. In this setting, the usual estimate of 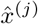 via total sum normalization will not suffice due to the preponderance of zero counts and the observed sequencing counts 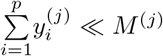, which induces severe artifacts in observed correlation due to differences in library size. For each row in the taxa table, we drew a single sample from the Dirichlet Multinomial model, given the input counts and the total size as the parameters for

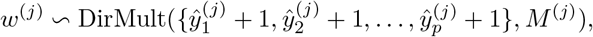

where the pseudo-count addition acts as a Laplace prior on the component proportions. We then supply total sum normalized simulated counts 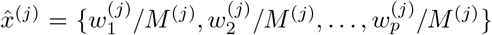 as the input data for the various network inference tools. We then collected our *n* count vectors into a data matrix *W* = {*w*^(1)^, *w*^(2)^, …, *w*^(*n*)^}^*T*^, and use log-transformed absolute covariances cov [log *W*], as the ground-truth matrices that need to be recovered by the inference schemes.

## American Gut Project data collection and high-quality reference

We acquired data from the American Gut Project (AGP) [37] from the qiita database [25]. We selected OTU tables from all 29 experimental rounds of AGP and filtered the combined dataset to *p* = 303 OTUs and *n* = 7722 samples. To assess the reproducibility of the different graphical models we construct a high-quality reference data set. We sorted the 29 rounds of the AGP by read count size, selected the top eight rounds (*n* = 3551), and concatenated the OTU tables.

We applied SE-slr, SE-glasso, SE-mb, SE-poisson, and SE-ising to the reference data set. For SE-slr, we fixed the rank of the matrix *L* to *r* = 42. From the remaining 21 AGP rounds we selected uniformly at random subsamples of size *n* ∈ {50, 3500} out of the total *n* = 4171 samples and applied each network inference method.

## Human gut microbiome data collection

We assembled a human gut microbiome data collection collection, that comprised 26 diverse datasets from the qiita database (https://qiita.ucsd.edu/).

All datasets were processed using the standard QIIME protocols, resulting in OTU tables in `biom` format - relevant here, as this is the end point of several different processing pipelines. For all datasets, OTUs were picked, and taxonomy assigned from untrimmed sequences using the sortmerna method under a 97% cluster similarity, and the closed-reference greengenes database. We performed all downstream data processing in R using the phyloseq package [77]. If necessary, we merged multiple `biom` files within a particular dataset (e.g. multiple experiments), keeping the set of common OTUs. We then did a first-pass filtering of the OTU table, keeping only fecal samples. Next, we pruned low abundant samples, defined as being in the bottom 10% of total counts for the dataset, and also filtered OTUs which were present in fewer than 37% of samples.

We also acquired sample metadata files, which are available in tabular text format from qiita. We “cleaned” these files to achieve consistent capitalization, abbreviation, data units, and nomenclature where possible. We also removed redundant categories, recorded all missing data to `NA`, and converted date/time stamps to a POSIX standard or factor representation. The data collection is available as part of our reproducible workflow (see Data Availablity statement below) and may be of independent interest for gut microbiome meta-analysis tasks.

### Edge distance

To compare the consistency between networks from different gut microbiome datasets, we restrict ourselves to sub-networks with common nodes, i.e., taxa. Formally, we are given 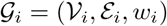 and 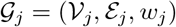 - two graphs inferred from datasets *i* and *j*, and with edge weights 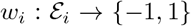 determined from the sign of the partial correlation. Let 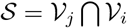, the set of nodes that have greengenes identifiers in common. Let 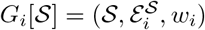 be the subgraph induced by this common node set, and denote by 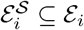 the subset of edges with endpoint vertices in 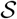. We then define the edge distance as

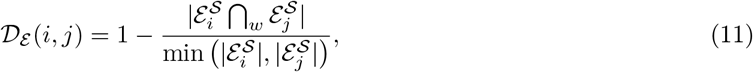

where ⋂_*w*_ denotes the sign-consistent intersection between edge sets and | · | is the size the set. The denominator ensures that the distance is scaled between 0 and 1.

For the meta-analysis, we required that each subgraph have at least 40 nodes and 50 edges, to avoid trivial comparisons between near-empty edge sets. This resulted in 215 pairwise comparisons across the 26 datasets.

### Consensus Gut Microbiome Network

For all networks inferred via SE-slr across the 26 gut microbiome datasets, we identified a consensus network from the learned node and edge sets. Using the greengenes identifier for each node, and the sign of partial correlations as edge weights, we merged sign-consistent edges that appeared in at least 5 individual datasets.

Additionally, we detected dense modules within the consensus network via the greedy clustering algorithm, implemented in the igraph package [68]. This resulted in 44 distinct network modules. We visualized the consensus SE-slr network in a force-directed layout, with OTUs colored by taxonomic order (Figure 4B).

### Data Availability

The code to reproduce methods and analysis in this work is available at https://github.com/zdk123/ SpiecEasiSLR_manuscript. The datasets, code, and containers with all required dependencies are available as a Synapse project: `syn20843558`.

## Appendix

### QIITA 16S rRNA sequencing datasets

**Table 1:** Key summary statistics of 16S rRNA sequencing datasets, downloaded from qiita at the study id[s] (third column). For each dataset, we assign a short name based on the study title (second column). We list the 16S variable `region` (forth column), sequencing platform (fifth column), number of samples *n* (sixth column) as well as number OTUs before and after filtering (column seven and eight, respectively). The last column lists the publications associated with the generation of the respective dataset (where available).

|  | name | qiita ids | region | platform | gut samples | OTUs | OTUs (filtered) | citation |
| --- | --- | --- | --- | --- | --- | --- | --- | --- |
| 1 | ABTX | 11052 | V4 | HiSeq | 241 | 17598 | 831 |  |
| 2 | AGP | 10317 | V4 | Multiple | 7722 | 1383 | 303 | [37] |
| 3 | Babies | 2010 | V4 | MiSeq | 132 | 11394 | 104 | [93] |
| 4 | Bangladesh | 10352 | V4 | HiSeq | 841 | 8229 | 796 | [107] |
| 5 | Bolivar | 11874 | V4 | MiSeq | 110 | 2331 | 476 | [101] |
| 6 | CCFA | 1939;1998 | V4 | MiSeq | 511 | 5326 | 452 | [95] |
| 7 | Colombian | 11993 | V4 | MiSeq | 396 | 6184 | 383 | [91, 92] |
| 8 | DHM | 11712 | V4 | HiSeq | 352 | 3277 | 101 |  |
| 9 | ECAM | 10249 | V4 | MiSeq | 963 | 1342 | 101 | [88] |
| 10 | Family | 797 | V2 | GAIIx | 184 | 15974 | 208 | [104] |
| 11 | Flores | 2151 | V4 | HiSeq | 777 | 9265 | 424 | [94] |
| 12 | GGMP | 11757 | V4 | HiSeq | 6308 | 16219 | 542 | [98] |
| 13 | Hadza | 11358 | V4 | MiSeq | 593 | 4783 | 597 | [102] |
| 14 | Herfarth | 2538 | V4 | HiSeq | 752 | 10530 | 456 | [106] |
| 15 | HIV | 11914 | V4 | MiSeq | 195 | 6771 | 439 | [86] |
| 16 | HMP | 1928 | V4 | 454 | 349 | 10730 | 189 | [90] |
| 17 | Infant | 10918 | V4 | MiSeq | 253 | 4682 | 170 |  |
| 18 | Malnourish | 1454 | V4 | HiSeq | 787 | 8680 | 274 | [105] |
| 19 | Mayo | 10184 | V4 | HiSeq | 865 | 6797 | 825 | [107] |
| 20 | Moving | 550 | V4 | GAIIx | 420 | 22765 | 537 | [89] |
| 21 | PNNL | 1629 | V4 | HiSeq | 614 | 10996 | 534 | [97] |
| 22 | PPR | 1774 | V4 | HiSeq | 122 | 20608 | 241 |  |
| 23 | Sandborn | 11546 | V4 | MiSeq | 292 | 4719 | 187 | [100] |
| 24 | Storage | 10394 | V4 | Multiple | 1263 | 9719 | 554 | [103] |
| 25 | TwinsUK | 2014 | V4 | MiSeq | 921 | 6842 | 767 | [96] |
| 26 | TZ | 2024 | V4 | MiSeq | 338 | 6868 | 294 | [87] |

### Robust PCA: top effect size

**Table 2:** List of metadata variables with largest *R*^2^ to low-rank components identified in the SE-slr model. The second column indicates the category membership of the variable, as described in the main text. The third column lists the variable name. The forth column denotes the number of datasets *n* the corresponding metadata variable was available in. The fifth column shows the best *R*^2^ between SE-slr low-rank components and the corresponding variable.

| | category | variable | n | $R^2$ |
| --- | --- | --- | --- | --- |
| 1 | III | zero_tax | 23 | 0.51 |
| 2 | I | host_subject_id | 10 | 0.87 |
| 3 | III | depth | 9 | 0.18 |
| 4 | I | age | 8 | 0.41 |
| 5 | I | sex | 7 | 0.17 |
| 6 | II | run_prefix | 6 | 0.40 |
| 7 | II | collection_timestamp | 4 | 0.38 |
| 8 | II | primer_plate | 4 | 0.29 |
| 9 | IV | diet | 3 | 0.50 |
| 10 | II | run_date | 3 | 0.28 |
| 11 | IV | antibiotics | 3 | 0.17 |
| 12 | II | extraction_robot | 3 | 0.10 |

### Identifiability of sparse and low rank inverse covariances from compositional data

We address the issue of identifiability for the estimator associated with the solutions of the convex program in (10). In particular, given estimates 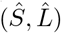 at fixed λ and *β*, we explore conditions under which the estimator can recover the pattern of non-zero entries, Supp(*A*) = {(*i*, *j*) : *A_i,j_* ≠ 0}, such that Supp (Ω^−1^) = Supp (*S*\*) and the rank(*L*\*) = 2, which are necessary prerequisites for exact recovery. For a general treatment of the problem, we refer to [110, 10]. Here, we focus on the particular problem of recovering sparse and low rank inverse covariances from compositional data without the presence of additional latent factors.

We build on results from [112] to obtain the deterministic identifiability conditions for sparse plus low rank recovery. We start by stating the following definition.

#### Definition 1

A matrix *S* is (*η*, *d*)-spread for *d* ∈ {1, …, *p*} and 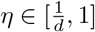 if and only if *d* is the maximum number of non-zero elements over the rows/columns of *S* (which is equivalent to the maximum degree+1 of the undirected graph) and

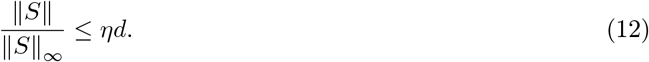

Here, ║·║ denotes the matrix operator norm (the maximum singular value), and ║·║_∞_ is the element-wise maximum magnitude of matrix elements. The (*η*, *d*)-spread condition implies that the underlying graph must be sparse and that edges must not be too concentrated on individual nodes, i.e., “hub nodes”.

#### Condition 1

The matrix *S*\* is (*η*, *d*)-spread.

#### Condition 2

Suppose *L*\* has the singular value decomposition 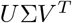, where *U*, 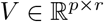 and 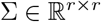 is a diagonal matrix. The singular vectors stored *U* and *V* must be (*r*, *μ*)-incoherent with respect to the standard basis. That is, for *r* ∈ {1, …, *p*} and 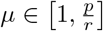 we require, for the symmetric *L*\*,

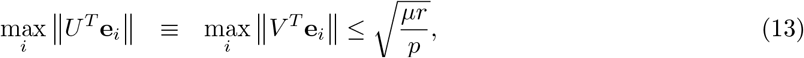

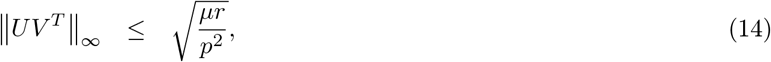

where **e**_*i*_’s are the standard basis vector and ║·║ is the vector ℓ^2^-norm. This condition requires that the singular vectors of the low-rank matrix are not too concentrated in a specific basis direction i.e. not too sparse.

#### Theorem 1 ([112])

Let Condition 1 and 2 hold. Then the matrix pair (*S*\*, *L*\*) is uniquely and deterministically identifiable via the estimator associated with (10) with appropriate tuning parameters if 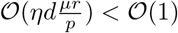.

**Proof of Theorem 1:** see [112]

#### Remark 1

Theorem 1 is a significant relaxation of the Theorem proposed in [110] which requires that *p* scales with *d*^2^*r*.

#### Remark 2

Assume we have an exact solution to (10) in form of the estimates 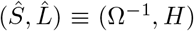. If the empirical covariance estimate is based on compositional data then 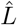 is not an arbitrary low-rank matrix but a function of Ω^−1^.

#### Lemma 1

Let 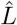 be a *p*-dimensional row vector. Let Ω^−1^*j* be a column vector of the row means of Ω^−1^. Let 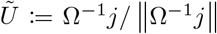, and let 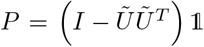 and 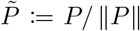 be column vectors and constructed such that 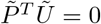. Then we have

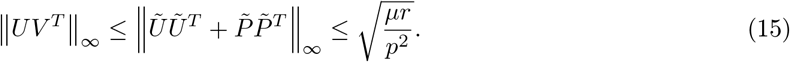

#### Remark 3

Before proving inequality (15), we discuss the consequences of having an SVD-free upper bound of the incoherence condition (14). The vector 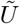 contains the (scaled) row means of the precision matrix, and *P* is the component of 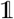 that is orthogonal to 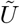. Therefore, each entry *i* in 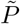 encodes the overall ‘agreement’ between the average precision of node *i* against all others. Therefore, coherence is satisfied when the scale of variance information for all variables is roughly equivalent. Assuming the upper bound is tight, overall incoherence is driven by outlier nodes.

This bound is reminiscent of an observation by Chayes [111], who described a special case of positive correlations between compositional variables being identifiable when the variances are identical. Thanks to a surprising “blessing of dimensionality”, the condition we describe is much less restrictive, since, if the matrix *S*\* is (*η*, *d*)-spread over many nodes, or, if positive and negative precisions are well balanced, then 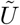 should tend toward a constant value.

#### Proof of Lemma 1

Let the matrix 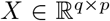 have rank *r* and the SVD 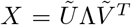, where 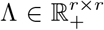 is the diagonal matrix with singular values along the diagonal. Let 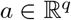 and 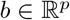 be arbitrary vectors. We are tasked with computing the SVD of

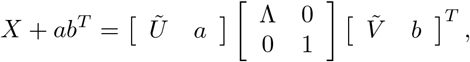

by modifications to 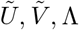.

Let 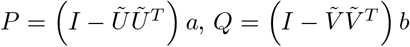 and let 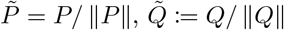. Then have that 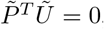, 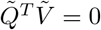 and

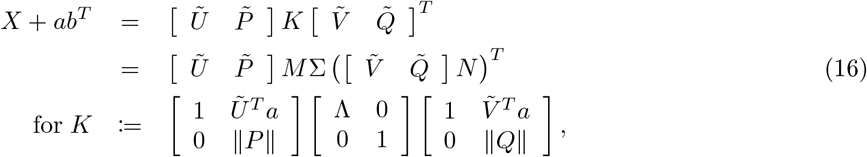

with *K* = *M*Σ*N^T^* as its SVD. 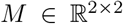 and 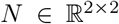 are rotations of the extended subspaces 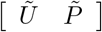 and 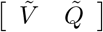, respectively.

Let (16) define a (modified Gram-Schmidt [108]) program that takes a previously computed SVD and new vectors as inputs and returns the updated SVD 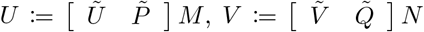 and diagonal matrix Σ.

Let *L*\* = *L*_1_ + *L*_2_ + *L*_3_ be a sum of three rank-1 matrices, *L*_1_ = Ω^−1^*J*, *L*_2_ = *J*Ω^−1^ and *L*_3_ = *J*Ω^−1^*J*. We can compute the SVD by recursively applying (16).

Let 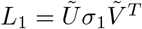. Since *L*_1_ is rank-1, we can define 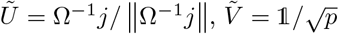 and 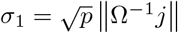, where 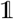 is the *p*-length all-ones vector and Ω^−1^*j* is the vector of row means of the precision matrix.

Now let *L*_2_ = *ab^T^*, with 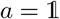 and *b* = Ω^−1^*j*. From (16), it holds that

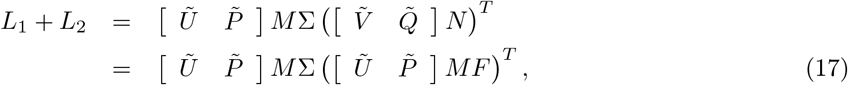

where 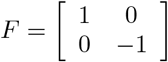 is a 90 degree reflection matrix. *L*_1_ + *L*_2_ is Hermitian so *L*_1_ + *L*_2_ = *UDU^T^*, where 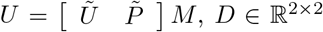 is the diagonal matrix of eigenvalues and |*D*| = Σ. We assert that *L*_1_ + *L*_2_ is indefinite and the second eigenvalue is negative by construction. Therefore *D* = Σ*F* which gives us the second line of equation 17.

Finally, let *L*_3_ = *ab^T^*, with *a* = *J*Ω^−1^*j* and 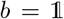. To complete the update of the SVD, application of (16) involves extending the (rotated) subspace 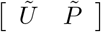. However, we have that both *a* and *b* are in the null space of 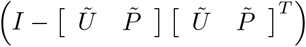. Therefore, the final update involves an additional rotation of 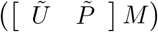. That is, the output of a second application of 17 gives us

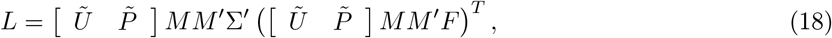

where 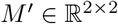 is the additional rotation matrix.

Let 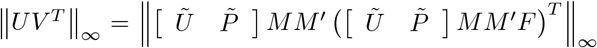. To find an SVD-free upper bound, we can compute the matrix 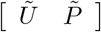 as above and solve

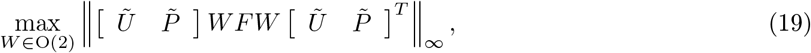

where 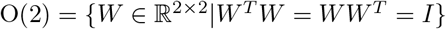 is the orthogonal group; distance-preserving transformations of a Euclidean space that also preserve the origin. To solve (19), we seek a rotation of 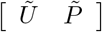 such that the longest component row vector is maximally oriented towards its basis direction in 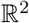. The index of this component vector is 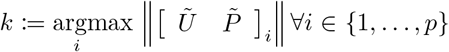, where [*A*]_*i*_ indicates the *i*th row vector of a matrix.

Let 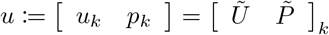 with 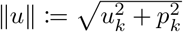. We can construct a Givens rotation matrix

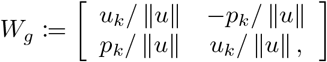

which is a (non-unique) solution for (19).

For any *i*, *j* ∈ {1, …, *p*} \ *k*, select component vectors 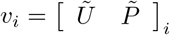 and 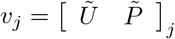. Recall that 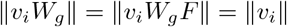, since the rotation is 2-norm invariant. Then we have

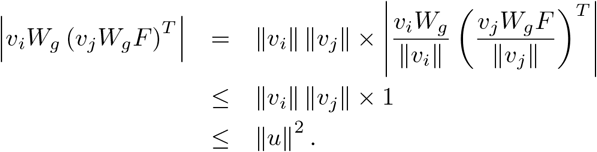

The last inequality comes from our definition of *u* as the ‘longest’ available vector. Finally, we have that

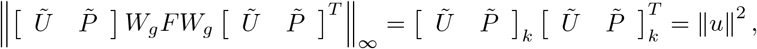

which concludes the proof.

## References

[1] John Aitchison. The statistical analysis of compositional data. Chapman and Hall, London; New York, 1986.

[2] Miklós Bálint, Mohammad Bahram, A. Murat Eren, Karoline Faust, Jed A. Fuhrman, Björn Lindahl, Robert B. O’Hara, Maarja Öpik, Mitchell L. Sogin, Martin Unterseher, and Leho Tedersoo. Millions of reads, thousands of taxa: microbial community structure and associations analyzed via marker genes. FEMS microbiology reviews, 6(24):189–96, 2016.

[3] Surojit Biswas, Meredith Mcdonald, Derek S Lundberg, Jeffery L. Dangl, and Vladimir Jojic. Learning microbial interaction networks from metagenomic count data. Journal of Computational Biology, 23(6):526–535, 2016.

[4] Johannes R. Björk, Francis K.C. Hui, Robert B. O’Hara, and Jose M. Montoya. Uncovering the drivers of host-associated microbiota with joint species distribution modelling. Molecular Ecology, 27(12):2714–2724, 2018.

[5] Ilana L. Brito, Thomas Gurry, Shijie Zhao, Katherine Huang, Sarah K. Young, Terrence P. Shea, Waisea Naisilisili, Aaron P. Jenkins, Stacy D. Jupiter, Dirk Gevers, and Eric J. Alm. Transmission of human-associated microbiota along family and social networks. Nature Microbiology, 4(6):964–971, 2019.

[6] Benjamin J. Callahan, Paul J. McMurdie, Michael J. Rosen, Andrew W. Han, Amy Jo A. Johnson, and Susan P. Holmes. DADA2: High-resolution sample inference from Illumina amplicon data. Nature Methods, 2016.

[7] Emmanuel J. Candès, Xiaodong Li, Yi Ma, and John Wright. Robust principal component analysis? Journal of the ACM, 2011.

[8] Yuanpei Cao, Wei Lin, and Hongzhe Li. Large covariance estimation for compositional data via composition-adjusted thresholding. Journal of the American Statistical Association, 114(526):759–772, 2019.

[9] Alex Carr, Christian Diener, Nitin S. Baliga, and Sean M. Gibbons. Use and abuse of correlation analyses in microbial ecology. ISME Journal, 2019.

[10] Venkat Chandrasekaran, Pablo A. Parrilo, and Alan S. Willsky. Latent variable graphical model selection via convex optimization. The Annals of Statistics, 40(4):1935–1967, 2012.

[11] I. Cho and M.J. Blaser. The human microbiome: at the interface of health and disease. Nature Reviews Genetics, 13(4):260–270, 2012.

[12] Ashley R. Coenen and Joshua S. Weitz. Limitations of Correlation-Based Inference in Complex Virus-Microbe Communities. mSystems, 3(4):7–9, 2018.

[13] Integrative HMP (iHMP) Research Network Consortium. The integrative human microbiome project: Dynamic analysis of microbiome-host omics profiles during periods of human health and disease, 2014.

[14] Lawrence A David, Corinne F Maurice, Rachel N Carmody, David B Gootenberg, Julie E Button, Benjamin E Wolfe, Alisha V Ling, A Sloan Devlin, Yug Varma, Michael A Fischbach, Sudha B Biddinger, Rachel J Dutton, and Peter J Turnbaugh. Diet rapidly and reproducibly alters the human gut microbiome. Nature, 505(7484):559–63, 2014.

[15] Anders B Dohlman and Xiling Shen. Mapping the microbial interactome: Statistical and experimental approaches for microbiome network inference. Experimental Biology and Medicine, 2019.

[16] Huaying Fang, Chengcheng Huang, Hongyu Zhao, and Minghua Deng. gCoda: Conditional Dependence Network Inference for Compositional Data. Journal of Computational Biology, 24(7):699–708, 2017.

[17] Karoline Faust and Jeroen Raes. Microbial interactions: from networks to models. Nature Reviews Microbiology, 10(8):538–550, 2012.

[18] Jerome Friedman, Trevor Hastie, and Rob Tibshirani. Regularization Paths for Generalized Linear Models via Coordinate Descent. Journal of statistical software, 2010.

[19] Jerome Friedman, Trevor Hastie, and Robert Tibshirani. Sparse inverse covariance estimation with the graphical lasso. Biostatistics (Oxford, England), 9(3):432–441, 2008.

[20] Jonathan Friedman and Eric J Alm. Inferring correlation networks from genomic survey data. PLoS computational biology, 8(9):e1002687, sep 2012.

[21] Dirk Gevers, Rob Knight, Joseph F. Petrosino, Katherine Huang, Amy L. McGuire, Bruce W. Birren, Karen E. Nelson, Owen White, Barbara A. Methé, and Curtis Huttenhower. The Human Microbiome Project: A Community Resource for the Healthy Human Microbiome. PLoS Biology, 10(8), 2012.

[22] Sean M Gibbons, Sean M Kearney, Chris S Smillie, and Eric J Alm. Two dynamic regimes in the human gut microbiome. PLoS Computational Biology, 13(2), 2017.

[23] Jack A Gilbert, Janet K Jansson, and Rob Knight. The Earth Microbiome project: successes and aspirations. BMC Biology, 2014.

[24] Gregory B. Gloor, Jia Rong Wu, Vera Pawlowsky-Glahn, and Juan José Egozcue. It’s all relative: analyzing microbiome data as compositions. Annals of Epidemiology, 2016.

[25] Antonio Gonzalez, Jose A. Navas-Molina, Tomasz Kosciolek, Daniel McDonald, Yoshiki Vázquez-Baeza, Gail Ackermann, Jeff DeReus, Stefan Janssen, Austin D. Swafford, Stephanie B. Orchanian, Jon G. Sanders, Joshua Shorenstein, Hannes Holste, Semar Petrus, Adam Robbins-Pianka, Colin J. Brislawn, Mingxun Wang, Jai Ram Rideout, Evan Bolyen, Matthew Dillon, J. Gregory Caporaso, Pieter C. Dorrestein, and Rob Knight. Qiita: rapid, web-enabled microbiome meta-analysis. Nature Methods, 15(10):796–798, 2018.

[26] Ethan T. Hillman, Hang Lu, Tianming Yao, and Cindy H. Nakatsu. Microbial Ecology along the Gastrointestinal Tract. Microbes and Environments, 32(4):300–313, 2017.

[27] Cho-jui Hsieh. QUIC: Quadratic Approximation for Sparse Inverse Covariance Estimation. Journal of Machine Learning Research, 15:2911–2947, 2014.

[28] Feng Ju and Tong Zhang. 16S rRNA gene high-throughput sequencing data mining of microbial diversity and interactions. Applied Microbiology and Biotechnology, pages 4119–4129, 2015.

[29] M. Senthil Kumar, Eric V. Slud, Kwame Okrah, Stephanie C. Hicks, Sridhar Hannenhalli, and Héctor Corrada Bravo. Analysis and correction of compositional bias in sparse sequencing count data. BMC genomics, 19(1):799, 2018.

[30] Zachary D Kurtz, Christian L Müller, Emily R Miraldi, Dan R Littman, Martin J Blaser, and Richard A Bonneau. Sparse and Compositionally Robust Inference of Microbial Ecological Networks. PLoS computational biology, 11(5):e1004226, may 2015.

[31] Christian L. Lauber, Micah Hamady, Rob Knight, and Noah Fierer. Pyrosequencing-based assessment of soil pH as a predictor of soil bacterial community structure at the continental scale. Applied and Environmental Microbiology, 2009.

[32] Gipsi Lima-Mendez, Karoline Faust, Nicolas Henry, Johan Decelle, Sébastien Colin, Fabrizio Carcillo, Samuel Chaffron, J. Cesar Ignacio-Espinosa, Simon Roux, Flora Vincent, Lucie Bittner, Youssef Darzi, Jun Wang, Stéphane Audic, Léo Berline, Gianluca Bontempi, Ana M. Cabello, Laurent Coppola, Francisco M. Cornejo-Castillo, Francesco D’Ovidio, Luc De Meester, Isabel Ferrera, Marie José Garet-Delmas, Lionel Guidi, Elena Lara, Stéphane Pesant, Marta Royo-Llonch, Guillem Salazar, Pablo Sánchez, Marta Sebastian, Caroline Souffreau, Céline Dimier, Marc Picheral, Sarah Searson, Stefanie Kandels-Lewis, Gabriel Gorsky, Fabrice Not, Hiroyuki Ogata, Sabrina Speich, Lars Stemmann, Jean Weissenbach, Patrick Wincker, Silvia G. Acinas, Shinichi Sunagawa, Peer Bork, Matthew B. Sullivan, Eric Karsenti, Chris Bowler, Colomban De Vargas, Jeroen Raes, Emmanuel Boss, Michael Follows, Nigel Grimsley, Pascal Hingamp, Daniele Iudicone, Olivier Jaillon, Lee Karp-Boss, Uros Krzic, Emmanuel G. Reynaud, Christian Sardet, Mike Sieracki, and Didier Velayoudon. Determinants of community structure in the global plankton interactome. Science, 2015.

[33] Han Liu, Fang Han, Ming Yuan, John Lafferty, and Larry Wasserman. High Dimensional Semiparametric Gaussian Copula Graphical Models. The Annals of Statistics, 40(4):34, 2012.

[34] Han Liu, Kathryn Roeder, and Larry Wasserman. Stability Approach to Regularization Selection (StARS) for High Dimensional Graphical Models. Advances in neural information processing systems, 24(2):1432–1440, 2010.

[35] Ramiro Logares, Shinichi Sunagawa, Guillem Salazar, Francisco M. Cornejo-Castillo, Isabel Ferrera, Hugo Sarmento, Pascal Hingamp, Hiroyuki Ogata, Colomban de Vargas, Gipsi Lima-Mendez, Jeroen Raes, Julie Poulain, Olivier Jaillon, Patrick Wincker, Stefanie Kandels-Lewis, Eric Karsenti, Peer Bork, and Silvia G. Acinas. Metagenomic 16S rDNA Illumina tags are a powerful alternative to amplicon sequencing to explore diversity and structure of microbial communities. Environmental Microbiology, 2014.

[36] Shiqian Ma, Lingzhou Xue, and Hui Zou. Alternating Direction Methods for Latent Variable Gaussian Graphical Model Selection. Neural computation, (2012):1–36, apr 2013.

[37] Daniel McDonald, Embriette Hyde, Justine W. Debelius, James T. Morton, Antonio Gonzalez, Gail Ackermann, Alexander A. Aksenov, Bahar Behsaz, Caitriona Brennan, Yingfeng Chen, Lindsay DeRight Goldasich, Pieter C. Dorrestein, Robert R. Dunn, Ashkaan K. Fahimipour, James Gaffney, Jack A. Gilbert, Grant Gogul, Jessica L. Green, Philip Hugenholtz, Greg Humphrey, Curtis Huttenhower, Matthew A. Jackson, Stefan Janssen, Dilip V. Jeste, Lingjing Jiang, Scott T. Kelley, Dan Knights, Tomasz Kosciolek, Joshua Ladau, Jeff Leach, Clarisse Marotz, Dmitry Meleshko, Alexey V. Melnik, Jessica L. Metcalf, Hosein Mohimani, Emmanuel Montassier, Jose Navas-Molina, Tanya T. Nguyen, Shyamal Peddada, Pavel Pevzner, Katherine S. Pollard, Gholamali Rahnavard, Adam Robbins-Pianka, Naseer Sangwan, Joshua Shorenstein, Larry Smarr, Se Jin Song, Timothy Spector, Austin D. Swafford, Varykina G. Thackray, Luke R. Thompson, Anupriya Tripathi, Yoshiki Vázquez-Baeza, Alison Vrbanac, Paul Wischmeyer, Elaine Wolfe, Qiyun Zhu, Rob Knight, Allison E. Mann, Amnon Amir, Angel Frazier, Cameron Martino, Carlito Lebrilla, Catherine Lozupone, Cecil M. Lewis, Charles Raison, Chi Zhang, Christian L. Lauber, Christina Warinner, Christopher A. Lowry, Chris Callewaert, Cinnamon Bloss, Dana Willner, Daniela Domingos Galzerani, David J. Gonzalez, David A. Mills, Deepak Chopra, Dirk Gevers, Donna Berg-Lyons, Dorothy D. Sears, Doug Wendel, Elijah Lovelace, Emily Pierce, Emily TerAvest, Evan Bolyen, Frederic D. Bushman, Gary D. Wu, George M. Church, Gordon Saxe, Hanna D. Holscher, Ivo Ugrina, J. Bruce German, J. Gregory Caporaso, Jacob M. Wozniak, Jacqueline Kerr, Jacques Ravel, James D. Lewis, Jan S. Suchodolski, Janet K. Jansson, Jarrad T. Hampton-Marcell, Jason Bobe, Jeroen Raes, John H. Chase, Jonathan A. Eisen, Jonathan Monk, Jose C. Clemente, Joseph Petrosino, Julia Goodrich, Julia Gauglitz, Julian Jacobs, Karsten Zengler, Kelly S. Swanson, Kim Lewis, Kris Mayer, Kyle Bittinger, Lindsay Dillon, Livia S. Zaramela, Lynn M. Schriml, Maria G. Dominguez-Bello, Marta M. Jankowska, Martin Blaser, Meg Pirrung, Michael Minson, Mike Kurisu, Nadim Ajami, Neil R. Gottel, Nicholas Chia, Noah Fierer, Owen White, Patrice D. Cani, Pawel Gajer, Philip Strandwitz, Purna Kashyap, Rachel Dutton, Rachel S. Park, Ramnik J. Xavier, Robert H. Mills, Rosa Krajmalnik-Brown, Ruth Ley, Sarah M. Owens, Scott Klemmer, Sébastien Matamoros, Siavash Mirarab, Stephanie Moorman, Susan Holmes, Tara Schwartz, Tifani W. Eshoo-Anton, Tim Vigers, Vineet Pandey, Will Van Treuren, Xin Fang, Zhenjiang Zech Xu, Alan Jarmusch, Justin Geier, Nicolai Reeve, Ricardo Silva, Evguenia Kopylova, Dominic Nguyen, Karenina Sanders, Rodolfo Antonio Salido Benitez, Arthur Cole Heale, Max Abramson, Jérôme Waldispühl, Alexander Butyaev, Chris Drogaris, Elena Nazarova, Madeleine Ball, and Beau Gunderson. American Gut: an Open Platform for Citizen Science Microbiome Research. mSystems, 2018.

[38] Michael R. McLaren, Amy D. Willis, and Benjamin J. Callahan. Consistent and correctable bias in metagenomic sequencing experiments. eLife, 2019.

[39] Paul J. McMurdie and Susan Holmes. Waste Not, Want Not: Why Rarefying Microbiome Data Is Inadmissible. PLoS Computational Biology, 10(4):e1003531, apr 2014.

[40] Nicolai Meinshausen and Peter Bühlmann. High Dimensional Graphs and Variable Selection with the Lasso. The Annals of Statistics, 34(3):1436–1462, 2006.

[41] Rajita Menon, Vivek Ramanan, and Kirill S. Korolev. Interactions between species introduce spurious associations in microbiome studies. PLoS Computational Biology, 14(1):1–20, 2018.

[42] Christian L. Müller, Richard Bonneau, and Zachary Kurtz. Generalized Stability Approach for Regularized Graphical Models. may 2016.

[43] Pradeep Ravikumar, Martin J. Wainwright, Garvesh Raskutti, and Bin Yu. High-dimensional covariance estimation by minimizing ℓ_1_-penalized log-determinant divergence. Electronic Journal of Statistics, 5(January 2010):935–980, 2011.

[44] Victoria E. Ruiz, Thomas Battaglia, Zachary D. Kurtz, Luc Bijnens, Amy Ou, Isak Engstrand, Xuhui Zheng, Tadasu Iizumi, Briana J. Mullins, Christian L. Müller, Ken Cadwell, Richard Bonneau, Guillermo I. Perez-Perez, and Martin J. Blaser. A single early-in-life macrolide course has lasting effects on murine microbial network topology and immunity. Nature Communications, 8(1), 2017.

[45] T. M. Schmidt, E F DeLong, and N R Pace. Analysis of a marine picoplankton community by 16S rRNA gene cloning and sequencing. Journal of Bacteriology, 173(14):4371–4378, 1991.

[46] Rashmi Sinha, Galeb Abu-Ali, Emily Vogtmann, Anthony A Fodor, Boyu Ren, Amnon Amir, Emma Schwager, Jonathan Crabtree, Siyuan Ma, Christian C Abnet, Rob Knight, Owen White, and Curtis Huttenhower. Assessment of variation in microbial community amplicon sequencing by the Microbiome Quality Control (MBQC) project consortium. Nature Biotechnology, 2017.

[47] Eric Smit, Paula Leeflang, Boet Glandorf, Jan Dirk Van Elsas, and Karel Wernars. Analysis of fungal diversity in the wheat rhizosphere by sequencing of cloned PCR-amplified genes encoding 18S rRNA and temperature gradient gel electrophoresis. Applied and Environmental Microbiology, 1999.

[48] Shinichi Sunagawa, Luis Pedro Coelho, Samuel Chaffron, Jens Roat Kultima, Karine Labadie, Guillem Salazar, Bardya Djahanschiri, Georg Zeller, Daniel R. Mende, Adriana Alberti, Francisco M. Cornejo-Castillo, Paul I. Costea, Corinne Cruaud, Francesco D’Ovidio, Stefan Engelen, Isabel Ferrera, Josep M. Gasol, Lionel Guidi, Falk Hildebrand, Florian Kokoszka, Cyrille Lepoivre, Gipsi Lima-Mendez, Julie Poulain, Bonnie T. Poulos, Marta Royo-Llonch, Hugo Sarmento, Sara Vieira-Silva, Céline Dimier, Marc Picheral, Sarah Searson, Stefanie Kandels-Lewis, Emmanuel Boss, Michael Follows, Lee Karp-Boss, Uros Krzic, Emmanuel G. Reynaud, Christian Sardet, Mike Sieracki, Didier Velayoudon, Chris Bowler, Colomban De Vargas, Gabriel Gorsky, Nigel Grimsley, Pascal Hingamp, Daniele Iudicone, Olivier Jaillon, Fabrice Not, Hiroyuki Ogata, Stephane Pesant, Sabrina Speich, Lars Stemmann, Matthew B. Sullivan, Jean Weissenbach, Patrick Wincker, Eric Karsenti, Jeroen Raes, Silvia G. Acinas, and Peer Bork. Structure and function of the global ocean microbiome. Science, 2015.

[49] Christine Y. Turenne, Steven E. Sanche, Daryl J. Hoban, James A. Karlowsky, and Amin M. Kabani. Rapid identification of fungi by using the ITS2 genetic region and an automated fluorescent capillary electrophoresis system. Journal of Clinical Microbiology, 1999.

[50] Corinne Vacher, Alireza Tamaddoni-Nezhad, Stefaniya Kamenova, Nathalie Peyrard, Yann Moalic, Régis Sabbadin, Loïc Schwaller, Julien Chiquet, M. Alex Smith, Jessica Vallance, Virgil Fievet, Boris Jakuschkin, and David A. Bohan. Learning Ecological Networks from Next-Generation Sequencing Data. In Advances in Ecological Research, volume 54, chapter One, pages 1–39. 2016.

[51] Doris Vandeputte, Gunter Kathagen, Kevin D’Hoe, Sara Vieira-Silva, Mireia Valles-Colomer, Joaõ Sabino, Jun Wang, Raul Y Tito, Lindsey De Commer, Youssef Darzi, Séverine Vermeire, Gwen Falony, and Jeroen Raes. Quantitative microbiome profiling links gut community variation to microbial load. Nature, 551(7681):507–511, 2017.

[52] Ying-wooi Wan, Genevera I Allen, Yulia Baker, Eunho Yang, Pradeep Ravikumar, Matthew Anderson, and Zhandong Liu. Open Access XMRF: an R package to fit Markov Networks to high-throughput genetics data. BMC Systems Biology, 10(Suppl 3), 2016.

[53] Yuqing Yang, Ning Chen, and Ting Chen. Inference of Environmental Factor-Microbe and Microbe-Microbe Associations from Metagenomic Data Using a Hierarchical Bayesian Statistical Model. Cell Systems, 4(1):129–137.e5, 2017.

[54] Grace Yoon, Irina Gaynanova, and Christian L. Müller. Microbial Networks in SPRING - Semiparametric Rank-Based Correlation and Partial Correlation Estimation for Quantitative Microbiome Data. Frontiers in Genetics, 10:1–33, 2019.

[55] Teng Zhang and Hui Zou. Sparse precision matrix estimation via lasso penalized D-trace loss. Biometrika, 101(1):103–120, 2014.

## References

[56] John Aitchison. The statistical analysis of compositional data. Chapman and Hall, London; New York, 1986.

[57] John Aitchison and Michael Greenacre. Biplots of compositional data. Journal of the Royal Statistical, 51(4):375–392, 2002.

[58] G.I. Allen and Z. Liu. A Log-Linear Graphical Model for Inferring Genetic Networks from High-Throughput Sequencing Data. IEEE International Conference on Bioinformatics and Biomedicine (BIBM), pages 1–19, 2012.

[59] Onureena Banerjee, Laurent El Ghaoui, and Alexandre D’Aspremont. Model selection through sparse maximum likelihood estimation for multivariate gaussian or binary data. The Journal of Machine …, 9:485–516, 2008.

[60] Surojit Biswas, Meredith Mcdonald, Derek S Lundberg, Jeffery L. Dangl, and Vladimir Jojic. Learning microbial interaction networks from metagenomic count data. Journal of Computational Biology, 23(6):526–535, 2016.

[61] E.J. Candes and B. Recht. Exact low-rank matrix completion via convex optimization. 2008 46th Annual Allerton Conference on Communication, Control, and Computing, (m):1–49, 2008.

[62] Emmanuel J. Candès, Xiaodong Li, Yi Ma, and John Wright. Robust principal component analysis? Journal of the ACM, 2011.

[63] Emmanuel J Candès and Benjamin Recht. Exact matrix completion via convex optimization. Foundations of Computational Mathematics, 9(6):717–772, 2009.

[64] Yuanpei Cao, Wei Lin, and Hongzhe Li. Large Covariance Estimation for Compositional Data via Composition-Adjusted Thresholding. 2016.

[65] J Gregory Caporaso, Justin Kuczynski, Jesse Stombaugh, Kyle Bittinger, Frederic D Bushman, Elizabeth K Costello, Noah Fierer, Antonio Gonzalez Peña, Julia K Goodrich, Jeffrey I Gordon, Gavin A Huttley, Scott T Kelley, Dan Knights, Jeremy E Koenig, Ruth E Ley, Catherine A Lozupone, Daniel McDonald, Brian D Muegge, Meg Pirrung, Jens Reeder, Joel R Sevinsky, Peter J Turnbaugh, William A Walters, Jeremy Widmann, Tanya Yatsunenko, Jesse Zaneveld, and Rob Knight. QIIME allows analysis of high-throughput community sequencing data. Nature methods, 7(5):335–6, may 2010.

[66] Venkat Chandrasekaran, Pablo A. Parrilo, and Alan S. Willsky. Latent variable graphical model selection via convex optimization. The Annals of Statistics, 40(4):1935–1967, 2012.

[67] Venkat Chandrasekaran, Sujay Sanghavi, Pablo A. Parrilo, and Alan S. Willsky. Rank-Sparsity Incoherence for Matrix Decomposition. SIAM Journal on Optimization, 21(2):572–596, 2009.

[68] Gabor Csardi and Tamas Nepusz. The igraph software package for complex network research. Inter-Journal, Complex Sy:1695, 2006.

[69] J J Egozcue. Isometric logratio transformations for compositional data analysis. Mathematical …, 35(3):279–300, 2003.

[70] Jerome Friedman, Trevor Hastie, and Robert Tibshirani. Sparse inverse covariance estimation with the graphical lasso. Biostatistics (Oxford, England), 9(3):432–441, 2008.

[71] Antonio Gonzalez, Jose A. Navas-Molina, Tomasz Kosciolek, Daniel McDonald, Yoshiki Vázquez-Baeza, Gail Ackermann, Jeff DeReus, Stefan Janssen, Austin D. Swafford, Stephanie B. Orchanian, Jon G. Sanders, Joshua Shorenstein, Hannes Holste, Semar Petrus, Adam Robbins-Pianka, Colin J. Brislawn, Mingxun Wang, Jai Ram Rideout, Evan Bolyen, Matthew Dillon, J. Gregory Caporaso, Pieter C. Dorrestein, and Rob Knight. Qiita: rapid, web-enabled microbiome meta-analysis. Nature Methods, 15(10):796–798, 2018.

[72] Ali Jalali and Sujay Sanghavi. Improved Deterministic Conditions for Sparse and Low-Rank Matrix Decomposition. 2016.

[73] Han Liu, Kathryn Roeder, and Larry Wasserman. Stability Approach to Regularization Selection (StARS) for High Dimensional Graphical Models. Advances in neural information processing systems, 24(2):1432–1440, 2010.

[74] Po-Ling Loh and Martin J Wainwright. Structure estimation for discrete graphical models: Generalized covariance matrices and their inverses. The Annals of Statistics, 41(6):3022–3049, 2013.

[75] Shiqian Ma, Lingzhou Xue, and Hui Zou. Alternating Direction Methods for Latent Variable Gaussian Graphical Model Selection. Neural computation, (2012):1–36, apr 2013.

[76] Daniel McDonald, Embriette Hyde, Justine W. Debelius, James T. Morton, Antonio Gonzalez, Gail Ackermann, Alexander A. Aksenov, Bahar Behsaz, Caitriona Brennan, Yingfeng Chen, Lindsay DeRight Goldasich, Pieter C. Dorrestein, Robert R. Dunn, Ashkaan K. Fahimipour, James Gaffney, Jack A. Gilbert, Grant Gogul, Jessica L. Green, Philip Hugenholtz, Greg Humphrey, Curtis Huttenhower, Matthew A. Jackson, Stefan Janssen, Dilip V. Jeste, Lingjing Jiang, Scott T. Kelley, Dan Knights, Tomasz Kosciolek, Joshua Ladau, Jeff Leach, Clarisse Marotz, Dmitry Meleshko, Alexey V. Melnik, Jessica L. Metcalf, Hosein Mohimani, Emmanuel Montassier, Jose Navas-Molina, Tanya T. Nguyen, Shyamal Peddada, Pavel Pevzner, Katherine S. Pollard, Gholamali Rahnavard, Adam Robbins-Pianka, Naseer Sangwan, Joshua Shorenstein, Larry Smarr, Se Jin Song, Timothy Spector, Austin D. Swafford, Varykina G. Thackray, Luke R. Thompson, Anupriya Tripathi, Yoshiki Vázquez-Baeza, Alison Vrbanac, Paul Wischmeyer, Elaine Wolfe, Qiyun Zhu, Rob Knight, Allison E. Mann, Amnon Amir, Angel Frazier, Cameron Martino, Carlito Lebrilla, Catherine Lozupone, Cecil M. Lewis, Charles Raison, Chi Zhang, Christian L. Lauber, Christina Warinner, Christopher A. Lowry, Chris Callewaert, Cinnamon Bloss, Dana Willner, Daniela Domingos Galzerani, David J. Gonzalez, David A. Mills, Deepak Chopra, Dirk Gevers, Donna Berg-Lyons, Dorothy D. Sears, Doug Wendel, Elijah Lovelace, Emily Pierce, Emily Ter-Avest, Evan Bolyen, Frederic D. Bushman, Gary D. Wu, George M. Church, Gordon Saxe, Hanna D. Holscher, Ivo Ugrina, J. Bruce German, J. Gregory Caporaso, Jacob M. Wozniak, Jacqueline Kerr, Jacques Ravel, James D. Lewis, Jan S. Suchodolski, Janet K. Jansson, Jarrad T. Hampton-Marcell, Jason Bobe, Jeroen Raes, John H. Chase, Jonathan A. Eisen, Jonathan Monk, Jose C. Clemente, Joseph Petrosino, Julia Goodrich, Julia Gauglitz, Julian Jacobs, Karsten Zengler, Kelly S. Swanson, Kim Lewis, Kris Mayer, Kyle Bittinger, Lindsay Dillon, Livia S. Zaramela, Lynn M. Schriml, Maria G. Dominguez-Bello, Marta M. Jankowska, Martin Blaser, Meg Pirrung, Michael Minson, Mike Kurisu, Nadim Ajami, Neil R. Gottel, Nicholas Chia, Noah Fierer, Owen White, Patrice D. Cani, Pawel Gajer, Philip Strandwitz, Purna Kashyap, Rachel Dutton, Rachel S. Park, Ramnik J. Xavier, Robert H. Mills, Rosa Krajmalnik-Brown, Ruth Ley, Sarah M. Owens, Scott Klemmer, Sébastien Matamoros, Siavash Mirarab, Stephanie Moorman, Susan Holmes, Tara Schwartz, Tifani W. Eshoo-Anton, Tim Vigers, Vineet Pandey, Will Van Treuren, Xin Fang, Zhenjiang Zech Xu, Alan Jarmusch, Justin Geier, Nicolai Reeve, Ricardo Silva, Evguenia Kopylova, Dominic Nguyen, Karenina Sanders, Rodolfo Antonio Salido Benitez, Arthur Cole Heale, Max Abramson, Jérôme Waldispühl, Alexander Butyaev, Chris Drogaris, Elena Nazarova, Madeleine Ball, and Beau Gunderson. American Gut: an Open Platform for Citizen Science Microbiome Research. mSystems, 2018.

[77] Paul J McMurdie and Susan Holmes. phyloseq: An R Package for Reproducible Interactive Analysis and Graphics of Microbiome Census Data. PloS one, 8(4):e61217, jan 2013.

[78] Paul J. McMurdie and Susan Holmes. Waste Not, Want Not: Why Rarefying Microbiome Data Is Inadmissible. PLoS Computational Biology, 10(4):e1003531, apr 2014.

[79] Nicolai Meinshausen and Peter Bühlmann. High Dimensional Graphs and Variable Selection with the Lasso. The Annals of Statistics, 34(3):1436–1462, 2006.

[80] Christian L. Müller, Richard Bonneau, and Zachary Kurtz. Generalized Stability Approach for Regularized Graphical Models. may 2016.

[81] Justin D Silverman, Alex D. Washburne, Sayan Mukherjee, and Lawrence A David. A phylogenetic transform enhances analysis of compositional microbiota data. eLife, 6:072413, aug 2017.

[82] Doris Vandeputte, Gunter Kathagen, Kevin D’Hoe, Sara Vieira-Silva, Mireia Valles-Colomer, Joaõ Sabino, Jun Wang, Raul Y Tito, Lindsey De Commer, Youssef Darzi, Séverine Vermeire, Gwen Falony, and Jeroen Raes. Quantitative microbiome profiling links gut community variation to microbial load. Nature, 551(7681):507–511, 2017.

[83] Ying-wooi Wan, Genevera I Allen, Yulia Baker, Eunho Yang, Pradeep Ravikumar, Matthew Anderson, and Zhandong Liu. Open Access XMRF : an R package to fit Markov Networks to high-throughput genetics data. BMC Systems Biology, 10(Suppl 3), 2016.

[84] Daniela M. Witten, Jerome H. Friedman, and Noah Simon. New Insights and Faster Computations for the Graphical Lasso. Journal of Computational and Graphical Statistics, 20(4):892–900, jan 2011.

[85] Eunho Yang, Pradeep Ravikumar, Genevera I Allen, and Zhandong Liu. On Graphical Models via Univariate Exponential Family Distributions. Journal of Machine Learning Research, 16:3813–3847, 2015.

## References

[86] Abigail J.S. Armstrong, Michael Shaffer, Nichole M. Nusbacher, Christine Griesmer, Suzanne Fiorillo, Jennifer M. Schneider, C. Preston Neff, Sam X. Li, Andrew P. Fontenot, Thomas Campbell, Brent E. Palmer, and Catherine A. Lozupone. An exploration of Prevotella-rich microbiomes in HIV and men who have sex with men. Microbiome, 2018.

[87] Jordan E. Bisanz, Megan K. Enos, George PrayGod, Shannon Seney, Jean M. Macklaim, Stephanie Chilton, Dana Willner, Rob Knight, Christoph Fusch, Gerhard Fusch, Gregory B. Gloor, Jeremy P. Burton, and Gregor Reid. Microbiota at multiple body sites during pregnancy in a rural tanzanian population and effects of Moringa-supplemented probiotic yogurt. Applied and Environmental Microbiology, 2015.

[88] Nicholas A. Bokulich, Jennifer Chung, Thomas Battaglia, Nora Henderson, Melanie Jay, Huilin Li, Arnon D. Lieber, Fen Wu, Guillermo I. Perez-Perez, Yu Chen, William Schweizer, Xuhui Zheng, Monica Contreras, Maria Gloria Dominguez-Bello, and Martin J. Blaser. Antibiotics, birth mode, and diet shape microbiome maturation during early life. Science Translational Medicine, 2016.

[89] J Gregory Caporaso, Christian L Lauber, Elizabeth K Costello, Donna Berg-Lyons, Antonio Gonzalez, Jesse Stombaugh, Dan Knights, Pawel Gajer, Jacques Ravel, Noah Fierer, Jeffrey I Gordon, and Rob Knight. Moving pictures of the human microbiome. Genome biology, 12(5):R50, jan 2011.

[90] The Human Microbiome Project Consortium. Structure, function and diversity of the healthy human microbiome. Nature, 486(7402):207–214, 2012.

[91] J. De La Cuesta-Zuluaga, V. Corrales-Agudelo, J. A. Carmona, J. M. Abad, and J. S. Escobar. Body size phenotypes comprehensively assess cardiometabolic risk and refine the association between obesity and gut microbiota. International Journal of Obesity, 2018.

[92] Jacobo De La Cuesta-Zuluaga, J, Vanessa Corrales-Agudelo, Eliana P. Velásquez-Mejía, Jenny A. Carmona, José M. Abad, and Juan S. Escobar. Gut microbiota is associated with obesity and cardiometabolic disease in a population in the midst of Westernization. Scientific Reports, 2018.

[93] Maria G. Dominguez-Bello, Kassandra M. De Jesus-Laboy, Nan Shen, Laura M. Cox, Amnon Amir, Antonio Gonzalez, Nicholas A. Bokulich, Se Jin Song, Marina Hoashi, Juana I. Rivera-Vinas, Keimari Mendez, Rob Knight, and Jose C. Clemente. Partial restoration of the microbiota of cesarean-born infants via vaginal microbial transfer. Nature Medicine, 2016.

[94] Gilberto E. Flores, J. Gregory Caporaso, Jessica B. Henley, Jai R.am Rideout, Daniel Domogala, John Chase, Jonathan W. Leff, Yoshiki Vázquez-Baeza, Antonio Gonzalez, Rob Knight, Robert R. Dunn, and Noah Fierer. Temporal variability is a personalized feature of the human microbiome. Genome biology, 2014.

[95] Dirk Gevers, Subra Kugathasan, Lee A Denson, Yoshiki Vázquez-Baeza, Will Van Treuren, Boyu Ren, Emma Schwager, Dan Knights, Se Jin Song, Moran Yassour, Xochitl C Morgan, Aleksandar D Kostic, Chengwei Luo, Antonio González, Daniel McDonald, Yael Haberman, Thomas Walters, Susan Baker, Joel Rosh, Michael Stephens, Melvin Heyman, James Markowitz, Robert Baldassano, Anne Griffiths, Francisco Sylvester, David Mack, Sandra Kim, Wallace Crandall, Jeffrey Hyams, Curtis Huttenhower, Rob Knight, and Ramnik J Xavier. The treatment-naive microbiome in new-onset Crohn’s disease. Cell host & microbe, 15(3):382–92, mar 2014.

[96] Julia K Goodrich, Jillian L Waters, Angela C Poole, Jessica L Sutter, Omry Koren, Ran Blekhman, Michelle Beaumont, William Van Treuren, Rob Knight, Jordana T Bell, Timothy D Spector, Andrew G Clark, and Ruth E Ley. Human genetics shape the gut microbiome. Cell, 159(4):789–799, nov 2014.

[97] Jonas Halfvarson, Colin J. Brislawn, Regina Lamendella, Yoshiki Vázquez-Baeza, William A. Walters, Lisa M. Bramer, Mauro D’Amato, Ferdinando Bonfiglio, Daniel McDonald, Antonio Gonzalez, Erin E. McClure, Mitchell F. Dunklebarger, Rob Knight, and Janet K. Jansson. Dynamics of the human gut microbiome in inflammatory bowel disease. Nature Microbiology, 2017.

[98] Yan He, Wei Wu, Hui Min Zheng, Pan Li, Daniel McDonald, Hua Fang Sheng, Mu Xuan Chen, Zi Hui Chen, Gui Yuan Ji, Zhong Dai Xi Zheng, Prabhakar Mujagond, Xiao Jiao Chen, Zu Hua Rong, Peng Chen, Li Yi Lyu, Xian Wang, Chong Bin Wu, Nan Yu, Yan Jun Xu, Jia Yin, Jeroen Raes, Rob Knight, Wen Jun Ma, and Hong Wei Zhou. Regional variation limits applications of healthy gut microbiome reference ranges and disease models, 2018.

[99] Daniel McDonald, Embriette Hyde, Justine W. Debelius, James T. Morton, Antonio Gonzalez, Gail Ackermann, Alexander A. Aksenov, Bahar Behsaz, Caitriona Brennan, Yingfeng Chen, Lindsay DeRight Goldasich, Pieter C. Dorrestein, Robert R. Dunn, Ashkaan K. Fahimipour, James Gaffney, Jack A. Gilbert, Grant Gogul, Jessica L. Green, Philip Hugenholtz, Greg Humphrey, Curtis Huttenhower, Matthew A. Jackson, Stefan Janssen, Dilip V. Jeste, Lingjing Jiang, Scott T. Kelley, Dan Knights, Tomasz Kosciolek, Joshua Ladau, Jeff Leach, Clarisse Marotz, Dmitry Meleshko, Alexey V. Melnik, Jessica L. Metcalf, Hosein Mohimani, Emmanuel Montassier, Jose Navas-Molina, Tanya T. Nguyen, Shyamal Peddada, Pavel Pevzner, Katherine S. Pollard, Gholamali Rahnavard, Adam Robbins-Pianka, Naseer Sangwan, Joshua Shorenstein, Larry Smarr, Se Jin Song, Timothy Spector, Austin D. Swafford, Varykina G. Thackray, Luke R. Thompson, Anupriya Tripathi, Yoshiki Vázquez-Baeza, Alison Vrbanac, Paul Wischmeyer, Elaine Wolfe, Qiyun Zhu, Rob Knight, Allison E. Mann, Amnon Amir, Angel Frazier, Cameron Martino, Carlito Lebrilla, Catherine Lozupone, Cecil M. Lewis, Charles Raison, Chi Zhang, Christian L. Lauber, Christina Warinner, Christopher A. Lowry, Chris Callewaert, Cinnamon Bloss, Dana Willner, Daniela Domingos Galzerani, David J. Gonzalez, David A. Mills, Deepak Chopra, Dirk Gevers, Donna Berg-Lyons, Dorothy D. Sears, Doug Wendel, Elijah Lovelace, Emily Pierce, Emily Ter-Avest, Evan Bolyen, Frederic D. Bushman, Gary D. Wu, George M. Church, Gordon Saxe, Hanna D. Holscher, Ivo Ugrina, J. Bruce German, J. Gregory Caporaso, Jacob M. Wozniak, Jacqueline Kerr, Jacques Ravel, James D. Lewis, Jan S. Suchodolski, Janet K. Jansson, Jarrad T. Hampton-Marcell, Jason Bobe, Jeroen Raes, John H. Chase, Jonathan A. Eisen, Jonathan Monk, Jose C. Clemente, Joseph Petrosino, Julia Goodrich, Julia Gauglitz, Julian Jacobs, Karsten Zengler, Kelly S. Swanson, Kim Lewis, Kris Mayer, Kyle Bittinger, Lindsay Dillon, Livia S. Zaramela, Lynn M. Schriml, Maria G. Dominguez-Bello, Marta M. Jankowska, Martin Blaser, Meg Pirrung, Michael Minson, Mike Kurisu, Nadim Ajami, Neil R. Gottel, Nicholas Chia, Noah Fierer, Owen White, Patrice D. Cani, Pawel Gajer, Philip Strandwitz, Purna Kashyap, Rachel Dutton, Rachel S. Park, Ramnik J. Xavier, Robert H. Mills, Rosa Krajmalnik-Brown, Ruth Ley, Sarah M. Owens, Scott Klemmer, Sébastien Matamoros, Siavash Mirarab, Stephanie Moorman, Susan Holmes, Tara Schwartz, Tifani W. Eshoo-Anton, Tim Vigers, Vineet Pandey, Will Van Treuren, Xin Fang, Zhenjiang Zech Xu, Alan Jarmusch, Justin Geier, Nicolai Reeve, Ricardo Silva, Evguenia Kopylova, Dominic Nguyen, Karenina Sanders, Rodolfo Antonio Salido Benitez, Arthur Cole Heale, Max Abramson, Jérôme Waldispühl, Alexander Butyaev, Chris Drogaris, Elena Nazarova, Madeleine Ball, and Beau Gunderson. American Gut: an Open Platform for Citizen Science Microbiome Research. mSystems, 2018.

[100] Ahmed Moustafa, Weizhong Li, Ericka L. Anderson, Emily H.M. Wong, Parambir S. Dulai, William J. Sandborn, William Biggs, Shibu Yooseph, Marcus B. Jones, J. Craig Venter, Karen E. Nelson, John T. Chang, Amalio Telenti, and Brigid S. Boland. Genetic risk, dysbiosis, and treatment stratification using host genome and gut microbiome in inflammatory bowel disease. Clinical and Translational Gastroenterology, 2018.

[101] Kelly V. Ruggles, Jincheng Wang, Angelina Volkova, Monica Contreras, Oscar Noya-Alarcon, Orlana Lander, Hortensia Caballero, and Maria G. Dominguez-Bello. Changes in the Gut Microbiota of Urban Subjects during an Immersion in the Traditional Diet and Lifestyle of a Rainforest Village. mSphere, 2018.

[102] Samuel A. Smits, Jeff Leach, Erica D. Sonnenburg, Carlos G. Gonzalez, Joshua S. Lichtman, Gregor Reid, Rob Knight, Alphaxard Manjurano, John Changalucha, Joshua E. Elias, Maria Gloria Dominguez-Bello, and Justin L. Sonnenburg. Seasonal cycling in the gut microbiome of the Hadza hunter-gatherers of Tanzania. Science, 2017.

[103] Se Jin Song, Amnon Amir, Jessica L. Metcalf, Katherine R. Amato, Zhenjiang Zech Xu, Greg Humphrey, and Rob Knight. Preservation Methods Differ in Fecal Microbiome Stability, Affecting Suitability for Field Studies. mSystems, 2016.

[104] Se Jin Song, Christian Lauber, Elizabeth K. Costello, Catherine A. Lozupone, Gregory Humphrey, Donna Berg-Lyons, J. Gregory Caporaso, Dan Knights, Jose C. Clemente, Sara Nakielny, Jeffrey I. Gordon, Noah Fierer, and Rob Knight. Cohabiting family members share microbiota with one another and with their dogs. eLife, 2013.

[105] Sathish Subramanian, Sayeeda Huq, Tanya Yatsunenko, Rashidul Haque, Mustafa Mahfuz, Mohammed A. Alam, Amber Benezra, Joseph Destefano, Martin F. Meier, Brian D. Muegge, Michael J. Barratt, Laura G. VanArendonk, Qunyuan Zhang, Michael A. Province, William A. Petri, Tahmeed Ahmed, and Jeffrey I. Gordon. Persistent gut microbiota immaturity in malnourished Bangladeshi children. Nature, 2014.

[106] Yoshiki Vázquez-Baeza, Antonio Gonzalez, Zhenjiang Zech Xu, Alex Washburne, Hans H. Herfarth, R. Balfour Sartor, and Rob Knight. Guiding longitudinal sampling in IBD cohorts, 2018.

[107] Emily Vogtmann, Jun Chen, Amnon Amir, Jianxin Shi, Christian C. Abnet, Heidi Nelson, Rob Knight, Nicholas Chia, Rashmi Sinha, Muhammad G. Kibriya, Yu Chen, Tariqul Islam, Mahbubul Eunes, Alauddin Ahmed, Jabun Naher, Anisur Rahman, Amnon Amir, Jianxin Shi, Christian C. Abnet, Heidi Nelson, Rob Knight, Nicholas Chia, Habibul Ahsan, and Rashmi Sinhaa. Comparison of fecal collection methods for microbiota studies in Bangladesh. Applied and Environmental Microbiology, 2017.

## References

[108] Matthew Brand. Fast low-rank modifications of the thin singular value decomposition. Linear Algebra and Its Applications, 415(1):20–30, may 2006.

[109] Venkat Chandrasekaran, Pablo A. Parrilo, and Alan S. Willsky. Latent variable graphical model selection via convex optimization. The Annals of Statistics, 40(4):1935–1967, 2012.

[110] Venkat Chandrasekaran, Sujay Sanghavi, Pablo A. Parrilo, and Alan S. Willsky. Rank-Sparsity Incoherence for Matrix Decomposition. SIAM Journal on Optimization, 21(2):572–596, 2009.

[111] F. Chayes. On Correlation between Variables of Constant Sum. Journal of Geophysical Research, 65(12):4185–4193, dec 1960.

[112] Ali Jalali and Sujay Sanghavi. Improved Deterministic Conditions for Sparse and Low-Rank Matrix Decomposition. 2016.

